# Structural basis for transcription initiation by bacterial ECF σ factors

**DOI:** 10.1101/534826

**Authors:** Lingting Li, Chengli Fang, Ningning Zhuang, Tiantian Wang, Yu Zhang

## Abstract

Bacterial RNA polymerase employs extra-cytoplasmic function (ECF) σ factors to regulate context-specific gene expression programs. Despite being the most abundant and divergent σ factor class, the structural basis of ECF σ factor-mediated transcription initiation remains unknown. Here, we determine a crystal structure of *Mycobacterium tuberculosis* (*Mtb*) RNAP holoenzyme comprising an RNAP core enzyme and the ECF σ factor σ^H^ (σ^H^-RNAP) at 2.7 Å, and solve another crystal structure of a transcription initiation complex of *Mtb* σ^H^-RNAP (σ^H^-RPo) comprising promoter DNA and an RNA primer at 2.8 Å. The two structures together reveal the interactions between σ^H^ and RNAP that are essential for σ^H^-RNAP holoenzyme assembly as well as the interactions between σ^H^-RNAP and promoter DNA responsible for stringent promoter recognition and for promoter unwinding. Our study establishes that ECF σ factors and primary σ factors employ distinct mechanisms for promoter recognition and for promoter unwinding.

## Introduction

Transcription initiation is the first and the most tightly regulated step of bacterial gene expression^1, 2, 3^. σ factors are required for transcription initiation^4^. After forming a complex with the RNA Polymerase (RNAP) core enzyme, σ factors guide RNAP to promoter DNA, open double-stranded DNA (dsDNA) to form a transcription bubble, facilitate synthesis of initial short RNA transcripts, and later assist in promoter escape^4, 5, 6^.

Bacterial σ factors are classified into two types—σ^70^- and σ^54^-type factors based on their distinct structures and mechanisms. The σ^70^-type factors can be further classified into four groups according to their conserved domains ^4^. Group-1 σ factors (or primary σ factors) contain domains σ_1.1_, σ_1.2_, σ_NCR_, σ_2_, σ_3.1_, σ_3.2_, σ_4_; group-2 σ factors contain all domains except σ_1.1_; group-3 σ factors contain σ_2_, σ_3.1_, σ_3.2_, σ_4_; while group-4 or ECF σ factors only contain σ_2_ and σ_4_^7^. The genomes of a majority of bacteria harbor one primary σ factor for expression of most genes (*i.e.*, group-1 σ factor; σ^70^ in *E. coli* and σ^A^ in gram-positive bacteria; referred as σ^A^ hereafter), and multiple alternative σ factors for expression of genes with cellular- or environmental-context-dependent functions ^8, 9^. The ECF σ factors are the largest family of alternative σ factors. On average, bacterial genomes encode six ECF σ factors; the given number for a particular bacterium will vary according to its genome size and environmental complexity^9, 10^. ECF σ factors enable bacteria to rapidly respond to a variety of stresses ^9, 11, 12^ and are known to be essential for the pathogenicity of several disease-causing bacteria ^13, 14^. *Mycobacterium tuberculosis* has 10 ECF σ factors (σ^C^, σ^D^, σ^E^, σ^G^, σ^H^, σ^I^, σ^J^, σ^K^, σ^L^, and σ^M^); deletion of ECF σ factors from *M. tuberculosis* results in attenuated disease progression (*e.g.*, *sigC* and *sigD*) or in alleviated virulence (*e.g.*, *sigE* and *sigH*) ^15, 16^.

The σ^A^ is capable of recognizing at least five conserved functional elements in the DNA sequences of gene promoters, including the “−35 element” (TTGACA)^17^, the “Z element” ^18^, the “extended −10 element” (TG)^19^, the “−10 element” (TATAAT) ^17^, and the “discriminator element” (GGG)^20^. Distinct domains of σ^A^ are responsible for interacting with these DNA elements: the domain σ_4_ forms sequence-specific interactions with exposed bases in the major groove of the −35 dsDNA ^21^; the σ_2.5_ and σ_3.1_ domains reach into the major groove of the extended −10 element and make base-specific contacts ^22, 23^; the σ_2_ and σ_1.2_ domains recognize and then unwind the −10 element dsDNA ^3, 24^.

During the process of promoter unwinding, a tryptophan dyad of σ_2_ (W256/W257 in *T. aquaticus* σ^A^ or W433/W434 in *E. coli* σ^70^) forms a chair-like structure that functions as a wedge to separate the dsDNA at the (−12)/(−11) junction ^22, 23^. The group-2 σ factors use the same set of residues to unwind promoter DNA; but the melting residues of group-3 σ factors are not conserved ^25^. Subsequently the base moieties of the unwound nucleotides at position −11 and −7 of the nontemplate strand–A_(−11)_(nt) and T_(−7)_(nt)–are flipped out and inserted into pre-formed pockets by σ_2_ and σ_1.2_ ^3, 24^. Domain σ_1.2_ also recognizes the discriminator element by flipping out the guanine base of G_(−6)_(nt) and inserting it into a pocket^24^. Although σ_3.2_ does not read the promoter sequence directly, it is essential for transcription initiation. Domain σ_3.2_ reaches into the RNAP active site cleft and “pre-organizes” template ssDNA^24^. Domain σ_3.2_ also blocks the path of the extending RNA chain (> 5nt) ^26, 27^ thereby contributing to both initial transcription pausing ^28^ and promoter escape ^29, 30^.

Each category of known ECF σ factors recognizes promoters bearing a unique sequence signature at the −35 and the −10 elements ^10, 31^. In contrast to the high tolerance to sequence variation at the −35 and the −10 promoter elements exhibited by the primary σ factor, the ECF σ factors have stringent requirements for sequence identity in the −35 and the −10 elements and for spacer length between these two elements through an unknown mechanism ^8, 32^. Although both the primary and ECF σ factors recognize the −35 element via σ_4_ and recognize the −10 element via σ_2_, the protein sequences of these two domains are not well conserved, and the consensus sequences of the two corresponding DNA elements vary. Crystal structures of individual σ_2_ or σ_4_ domains of ECF σ factors complexed with cognate DNA have suggested that these ECF domains bind the −35 and the −10 elements differently than does the primary σ factor, implicating a unique means of promoter recognition by ECF σ factors ^33, 34^. Another striking difference was revealed by a sequence analysis showing the surprising fact that ECF σ factors do not contain σ_3_ domains (σ_3.1_ and σ_3.2_), but instead contains a linker—highly variable in both length and sequence—to connect the σ_2_ and σ_4_ domains ^4^. This fact immediately raises the question of how these σ factors perform multiple steps of transcription initiation that the σ_3_ domain performs in the primary σ factor.

A recent crystal structure of *E. coli* σ^E^_2_/−10 ssDNA binary complex suggests that bacterial ECF σ factors probably recognize and unwind promoters through a unique mechanism. Specifically, *E. coli* σ^E^ employs a flexible ‘specificity loop’ to recognize a flipped master nucleotide of the −10 element and probably unwinds at a distinct position compared with that of σ^70^ by using non-conserved melting residues^34^. In contrast to the large collection of structural information of primary σ factor, no structure of bacterial RNAP complex with ECF σ factor is available. Therefore, it is largely unknown how ECF σ factors form a holoenzyme with RNAP and how ECF σ factors work alongside RNAP to recognize and to unwind promoter DNA. Here, we report the crystal structure of an ECF σ factor-RNAP holoenzyme comprising *M. tuberculosis* RNAP and σ^H^ at 2.70 Å resolution. We also report the crystal structure of an ECF σ factor-RNAP transcription initiation complex comprising *M. tuberculosis* σ^H^-RNAP holoenzyme, a full transcription bubble of promoter DNA, and an RNA primer at 2.80 Å resolution. The crystal structures present detailed interactions among RNAP, ECF σ factors, and promoter DNA. The structures together with data from biochemical assays collectively establish the structural basis of RNAP holoenzyme assembly, promoter recognition, and promoter unwinding by the ECF σ factors.

## Results

### The crystal structure of *M. tuberculosis* σ^H^-RNAP holoenzyme

The crystals of *M. tuberculosis* σ^H^-RNAP holoenzyme were unexpectedly obtained during an initial attempt to crystallize *M. tuberculosis* σ^H^-RPo (Supplementary Figure 1A, 1C-F). The crystal structure of *M. tuberculosis* σ^H^-RNAP holoenzyme at 2.7 Å resolution was determined by molecular replacement using a *M. smegmatis* RNAP core enzyme (PDB:5TW1) [https://www.rcsb.org/structure/5TW1] as the searching model ^35^. The Fo-Fc map shows unambiguous density for σ^H^ residues 22-195 (Table 1; Supplementary Figure 2A) and the anomalous difference map shows clear density for 4 out of 5 Se atoms, validating the σ^H^ model (Supplementary Figure 2A).

**TABLE 1.**
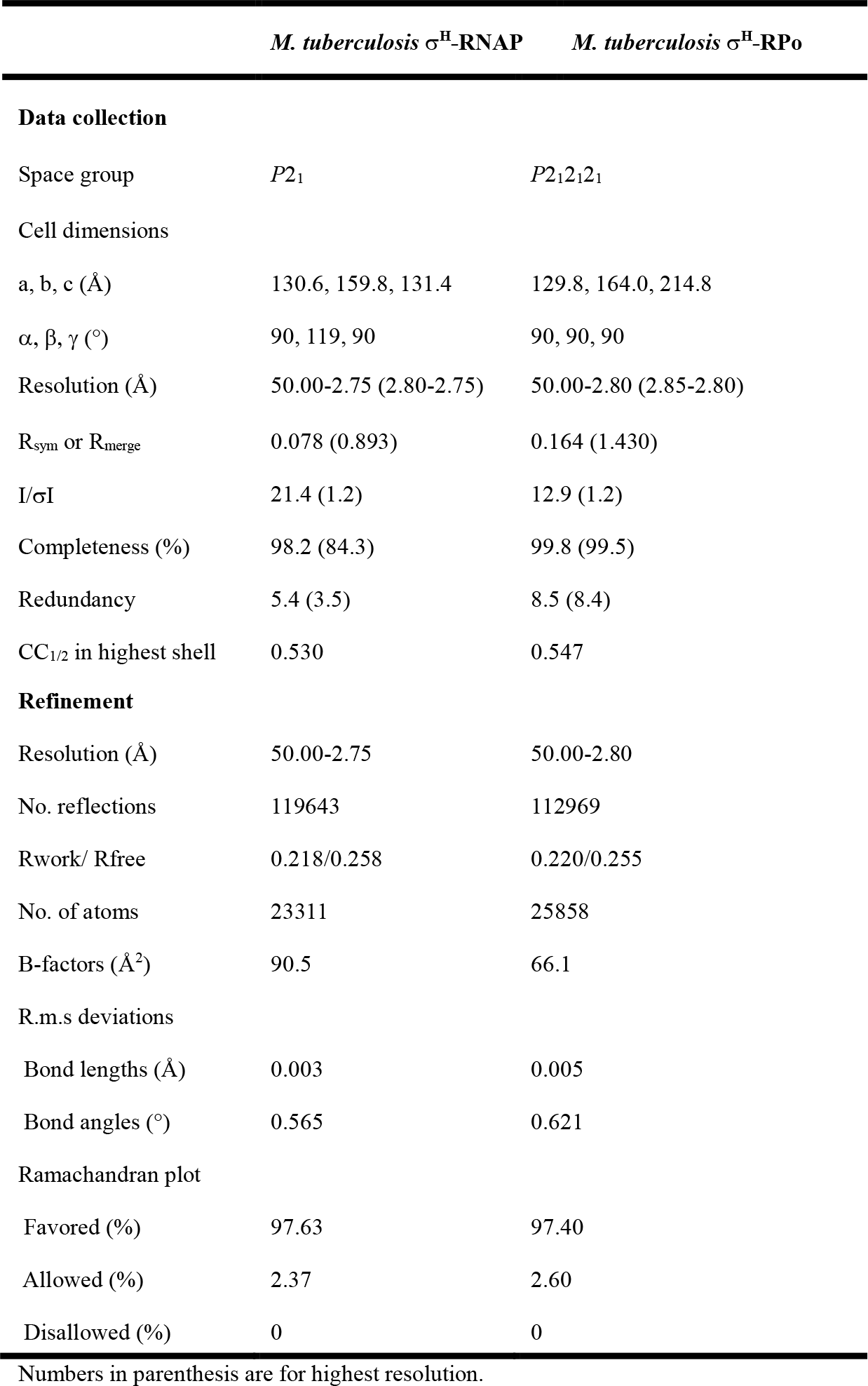
The statistics of crystal structures.

σ^H^_2_ (residues 22-99) and σ^H^_4_ (residues 140-195) fold into independent helical domains (Figure 1A and 1C). Lacking the σ_1.1_, σ_1.2_, and σ_NCR_ domains of σ^A^, the σ^H^_2_ domain is very compact, containing only four α helices (Figure 1C). The “specificity loop” (residues 72-79 in σ^H^_2_; Supplementary Figure 2A) known to be essential for recognition of the −10 element is disordered (no electron density), in contrast to the pre-organized specificity loop in the σ^A^-RNAP holoenzyme (Figure 1B, 1D and Supplementary Figure 2B-C). Lacking σ_1.2_, the domain that forms extensive interactions with the specificity loop of σ^A^, probably accounts for the disordered conformation of the specificity loop in σ^H^ (Figure 1C-D). As occurs in σ^A^_2_, the σ^H^_2_ domain resides in a cleft between the RNAP-β lobe and the RNAP-β′ coiled-coil (β′CC) and makes extensive electrostatic interactions with the latter (Figure 1E). Notably, the residues contacting β′CC of both σ^A^ and ECF σ factors are conserved (Supplementary Figure 2D-E and 3), suggesting that β′CC probably serves as an anchor point for the σ_2_ domain of the majority of bacterial σ factors.

**Figure 1.**
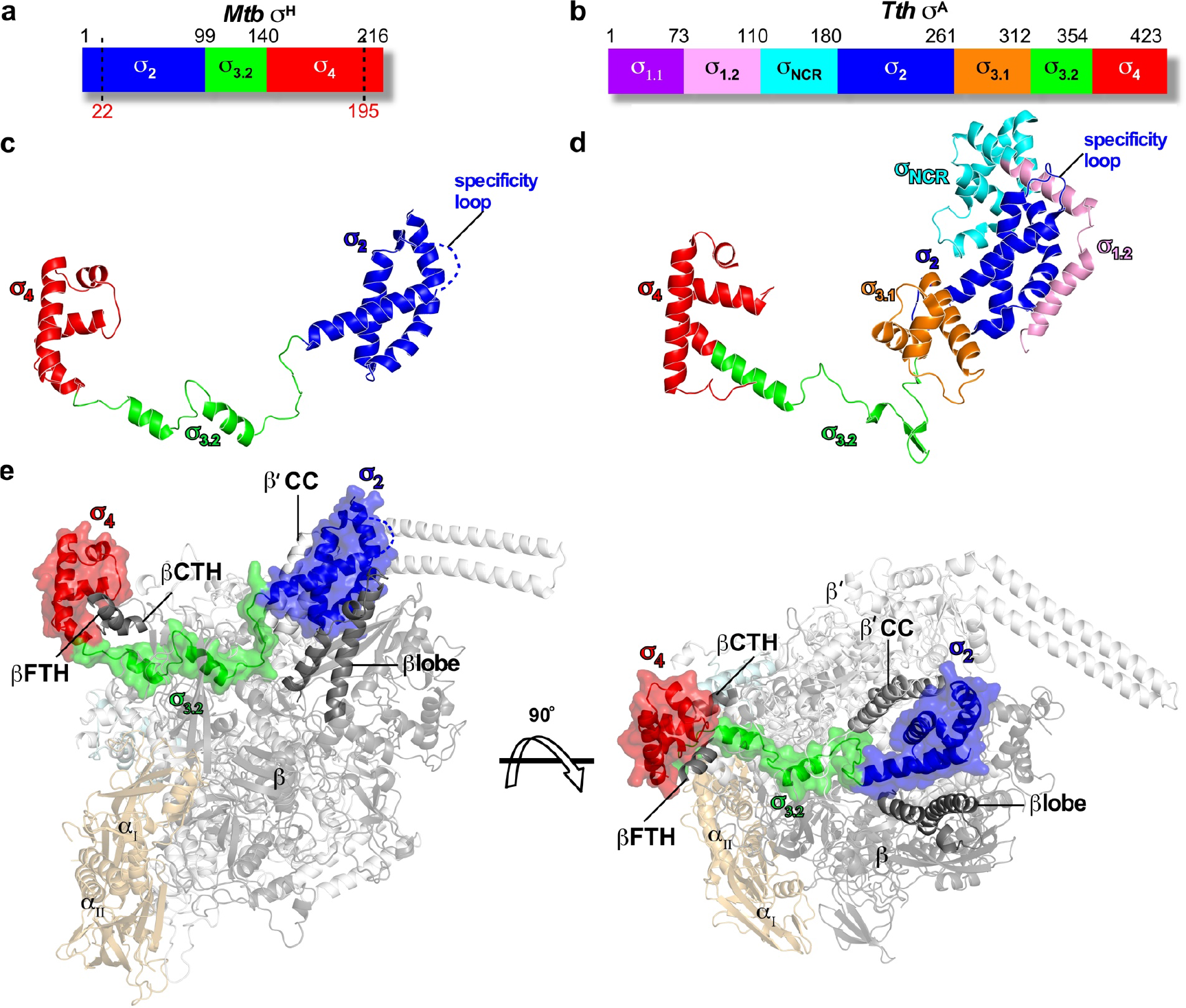
The crystal structure of *Mtb* σ^H^-RNAP holoenzyme. **(A)**Schematic diagram of *Mtb* σ^H^. Ordered regions in the structure are indicated by dashes. **(B)**Schematic diagram of *T. thermophiles* σ^A^. **(C)** σ^H^ in the crystal structure of *Mtb* σ^H^-RNAP holoenzyme. The disordered specificity loop is shown by blue dashes. **(D)** σ^A^ in the crystal structure of *T. thermophiles* σ^A^-RNAP (PDB: 1IW7) [https://www.rcsb.org/structure/1iw7] was used for comparison due to no available structure of *Mtb* σ^A^-RNAP holoenzyme. **(E)**Front and top views of *Mtb* σ^H^-RNAP holoenzyme. βFTH, the flap-tip helix on RNAP-βsubunit; βCTH, the C-terminal helix on RNAP-βsubunit; β’CC, the coiled-coil on RNAP-β’ subunit. σ^A^_1.1_, purple; σ^A^_1.2_, pink; σ^A^_NCR_, cyan; σ^2^, blue; σ^A^_3.1_, orange; σ_3.2_, green; σ_4_, red. RNAP-βsubunits, light orange; RNAP-βsubunit, black; RNAP-β′ subunit, gray; RNAP-ω subunit, light cyan.

The σ^H^_4_ domain enfolds the flap-tip helix of the RNAP-β subunit (βFTH; Figure 1E and 2A). The hydrophobic residues contacting the βFTH are conserved between the σ^A^ and the ECF σ factors (Supplementary Figure 2F-G and 3). Surprisingly, we discovered another anchor point for the σ^H^_4_ domain on RNAP—a C-terminal helix of the RNAP-β subunit (βCTH; residues 1145-1157; Figure 1E, 2A, and Supplementary Figure 2H and 2M). The interaction with βCTH was not observed in any of the previously reported bacterial σ^A^-RNAP structures^22, 24, 36, 37^. To explore the contribution of such interaction to the transcription activity of σ^H^-RNAP, we performed *in vitro* transcription experiments using wild-type or βCTH-deleted *Mtb* σ^H^-RNAP holoenzyme and *pClpB* promoter variants with −35/−10 spacer lengths ranging from 15-19 bp. The wild-type σ^H^-RNAP was most transcriptionally active with a promoter of 17-bp spacer (Figure 2B), consistent with a study reporting that most σ^H^-regulated promoters have a 17-bp spacer ^38^. The βCTH-deletion variant caused impaired transcription activity from promoter with the optimal spacer length (17 bp) but showed little effect on promoter with sub-optimal spacer lengths (16 bp and 18 bp) (Figure 2B), suggesting that the interactions between βCTH and σ^H^ are important for the transcription activity of σ^H^. Intriguingly, deletion of βCTH caused a general increase of σ^A^-dependent transcription activity from promoter with spacer lengths 15-19 bp (Supplementary Figure 2I).

**Figure 2.**
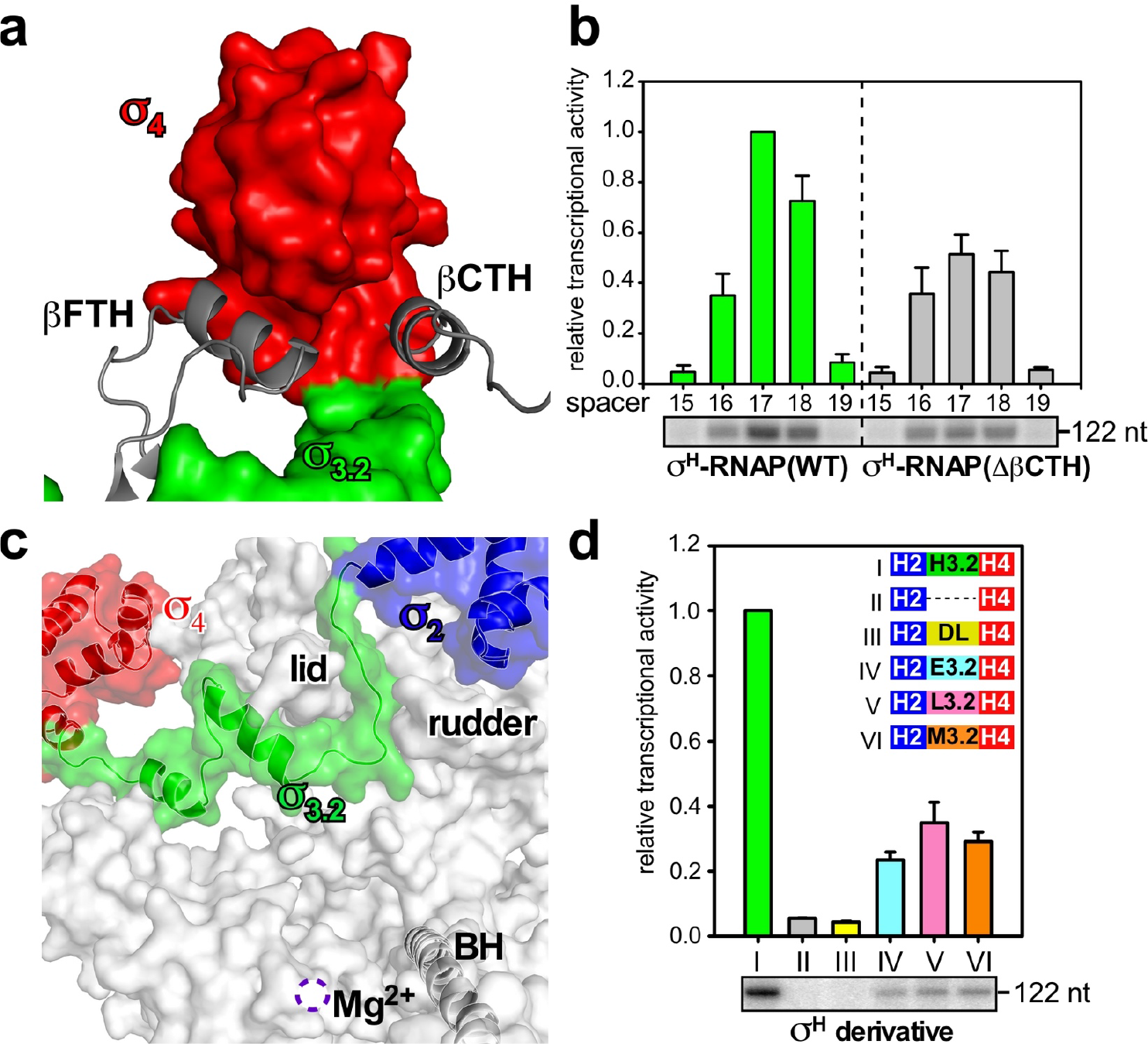
The interaction between *Mtb* RNAP core enzyme and σ^H^. (A) Both βFTH and βCTH interact with σ^H^_4_. Colors are as in previous figures. **(B)** The *in vitro* transcription activity of σ^H^-RNAP(WT) (green bars) and σ^H^-RNAP(ΔβCTH) (gray bars) from *pClpB* promoter variants with −35/−10 spacer length of 15-19 base pairs. “122 nt” indicates length of run-off RNA products. **(C)** The interaction between σ^H^_2_/σ^H^_4_ linker (σ^H^_3.2_) and RNAP core enzyme. RNAP-βsubunit was omitted for clarity. The location of catalytic Mg^2+^ is shown by a dashed purple circle. BH, bridge helix. RNAP-β′ subunit, gray. σ^H^_2_, σ^H^3.2, and σ^H^_4_, blue, green, and red respectively. **(D)**The *in vitro* transcription activity from *pClpB* promoter of RNAP holoenzymes comprising σ^H^ derivatives. H2-H3.2-H4, wild-type σ^H^; H2---H4, two individual domains of σ^H^_2_ and σ^H^_4_; H2-DL-H4, a chimeric σ^H^ with σ^H^_3.2_ replaced by a disordered loop with an equivalent residue number; H2-E3.2-H4, a chimeric σ^H^ with σ^H^_3.2_ replaced by *Mtb* σ^E^_3.2_; H2-L3.2-H4, a chimeric σ^H^ with σ^H^_3.2_ replaced by *Mtb* σ^L^_3.2_; H2-M3.2-H4, a chimeric σ^H^ with σ^H^_3.2_ replaced by *Mtb* σ^H^_3.2_. The experiments were repeated in triplicate, and the data are presented as mean ± S.E.M. Source data of (B) and (C) are provided as a Source Data file.

The most surprising finding in the structure of σ^H^-RNAP is the interaction between RNAP and the linker region connecting σ^H^_2_ and σ^H^_4_. The linker is the least conserved region among the ECF σ factors and shares no sequence similarity with the linker of σ^A^ (Supplementary Figure 3). Our structure shows that the linker region of σ^H^ dives into the active site cleft and emerges out from the RNA exit channel of RNAP (Figure 1E, 2C, and Supplementary Figure 4A-B). This interaction creates an entry channel for template ssDNA loading into the active site cleft during RPo formation, but blocks the exit pathway of extending RNA during subsequent transcription initiation events. The path of the σ^H^_2_/σ^H^_4_ linker in RNAP is similar to that of σ^H^_3.2_ (Supplementary Figure 4C-D), so we designated the σ^H^_2_/σ^H^_4_ linker as σ^H^_3.2_

To examine the significance of the interaction between σ^H^_3.2_ and RNAP, we tested the *in vitro* transcription activity of RNAP holoenzyme comprising σ^H^ variants with the σ^H^_3.2_ domain either deleted or swapped. Deleting σ^H^_3.2_ or replacing σ^H^_3.2_ with a protein sequence known to be disordered completely abolished the transcription activity of σ^H^ (Lanes II and III in Figure 2D), indicating that σ^H^_3.2_ does not simply serve as a σ^H^_2_/σ^H^_4_ linker; rather, the interaction between σ^H^_3.2_ and the active site cleft of RNAP is essential for transcription. Interestingly, replacing σ^H^_3.2_ with the σ_2_/σ_4_ linker from other ECF σ factor partially recovered the transcription activity (Lane IV, V, and VI in Figure 2D and Supplementary Figure 4E). Based on these results, we infer that other ECF σ factors probably have a functional σ_3.2_ domain that, while divergent in sequence, likely binds RNAP in a way somehow analogous to σ^H^_3.2_.

### The overall structure of *Mtb* σ^H^-RPo

To understand how σ^H^-RNAP holoenzyme recognizes promoter DNA and initiates transcription, we sought to determine a crystal structure of σ^H^-RPo. We assembled the complex by incubating the RNAP core enzyme, σ^H^, and a synthetic nucleic acid scaffold (Figure 3D, and Supplementary Figure 1B, 1G-J). The synthetic scaffold comprises an upstream DNA duplex (−34 to −10 with respect to transcription start site at +1) with a consensus −35 element (GGAACA), a non-complimentary transcription bubble (−9 to +2) with a consensus −10 element (GTT), a 7 nt RNA primer complimentary to template DNA (−6 to +1), and a downstream DNA duplex (+3 to +13). We determined the crystal structure of σ^H^-RPo at 2.8 Å resolution by molecular replacement using the crystal structure of *Mtb* σ^H^-RNAP holoenzyme as a search model (Table 1). The Fo-Fc map contoured at 2.5 σ shows clear density for all nucleotides of nontemplate ssDNA, template ssDNA, and RNA primer of the transcription bubble, as well as for all the nucleotides of the upstream and downstream DNA duplexes (Figure 3E).

**Figure 3.**
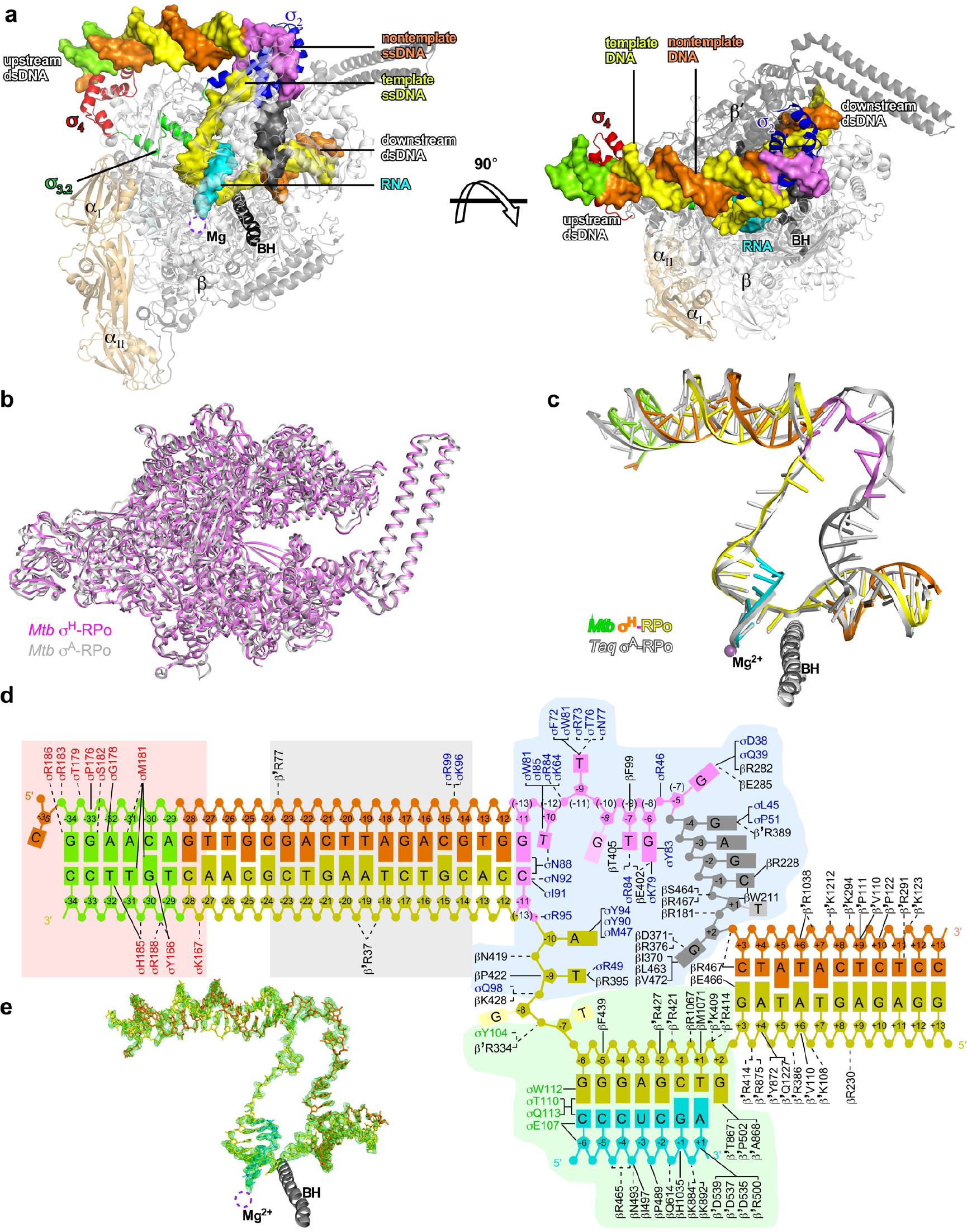
The crystal structure of *Mtb* σ^H^-RPo. **(A)** Front and top views of σ^H^-RPo overall structure. The α, β, β′, and ωsubunits of RNAP core enzyme are shown as ribbon and colored in light orange, gray, black, and light cyan respectively. The σ^H^_2_, σ^H^_3.2_ and σ^H^_4_ are shown as ribbon and colored as in above figures. The nontemplate DNA, template strand DNA, and RNA strands are shown in surface and colored in orange, yellow, and cyan, respectively, except the −35 element (green), the −10 element (purple), and the CRE (black). The location of the catalytic Mg^2+^ is indicated by a dashed circle. **(B)** Both *Mtb* σ^H^-RPo (violet) and *Mtb* σ^A^-RPo (light gray; PDB: 5UHA) [https://www.rcsb.org/structure/5uha] show closed clamp conformation. **(C)** The comparison of upstream dsDNA, transcription bubble and downstream dsDNA in *Mtb* σ^H^-RPo (colored as in figure 3A) and *Taq* σ^A^-RPo (light gray; PDB: 4XLN) [https://www.rcsb.org/structure/4XLN]. **(D)** Summary of protein-nucleic acid interactions. Solid line, van del Waals interactions; dashed line, polar interactions. Colors are as in above. Red box, interactions with the −35 element (details in Fig. 4A); gray box, interactions with the −35/−10 spacer (details in Fig. 4B); blue box, interactions of the ssDNA in transcription bubble (details in Fig. 4E, 5A-5D); green box, interactions with the DNA/RNA hybrid (details in Fig. 5E). The numbers in parenthesis are corresponding positions in σ^A^-regulated promoters. **(E)** The simulated-annealing omit Fo-Fc electron density map (nucleic-acids removed; green; contoured at 2.5 σ) and model for nucleic acids.

In the σ^H^-RPo structure, σ^H^ makes the same interactions with RNAP as in the structure of σ^H^-RNAP holoenzyme. The RNAP clamp adopts a closed conformation as in σ^A^-RPo ^24, 39^, consistent with previous smFRET results ^40^ and supporting the idea that clamp closure is also an obligatory step of RPo formation in ECF σ -mediated transcription initiation (Figure 3B).

The DNA/RNA hybrid resides in the active site cleft in a post-translocation state, and the downstream DNA duplex is accommodated in the main channel. The conformations of the DNA/RNA hybrid and downstream DNA duplex in σ^H^-RPo and σ^A^-RPo are similar (Figure 3C)^24^.

The σ^H^-RPo structure revealed multiple interactions responsible for promoter recognition and promoter unwinding by σ^H^-RNAP that we will describe in detail in each of the subsequent sections of our manuscript. These include: 1) σ^H^_4_ inserts into the major groove and reads the sequence of the −35 element (Figure 3A, 4A, and Supplementary Figure 6A); 2) the RNAP-β′ subunit stabilizes the upstream DNA duplex by contacting the phosphate backbone of nucleotides at positions −23, −20, and −19 (Figure 3A and 4B); 3) σ^H^_2_ unwinds dsDNA using an apparently distinct mechanism (Figure 3A and 4E); 4) σ^H^_2_ and the RNAP-β subunit recognize sequences at four positions in the −10 element via interactions with nontemplate ssDNA (Figure 3A and 5A-C); 5) the RNAP-β subunit recognizes the “CRE element” DNA sequence via interactions with nontemplate ssDNA (Figure 3A, 5D, and Supplementary Figure 7A); and 6) σ^H^_3.2_ guides the template ssDNA into the RNAP active center and forms interactions with the DNA/RNA hybrid in the active site cleft (Figure 3A, 5E, and Supplementary Figure 7G).

### The interactions of σ^H^_4_ with the −35 element

σ^H^-regulated promoters have a distinct consensus sequence at their −35 elements (5′-GGAAYA-3′; from −34 to −29; Supplementary Figure 5A) ^38, 40, 41^. Alternation of DNA sequences at each of the positions from −34 to −29 resulted in substantial loss of transcription activity (Supplementary Figure 5B). In the structure, σ^H^_4_ binds to the major groove of dsDNA of the −35 element and makes base-specific polar interactions with nucleotides at three (−34, −33, and −31) out of six positions (Figure 3D, 4A, and Supplementary 6A). The G_−34_(nt) makes two H-bonds with R186 through its O6 and N7 atoms; the G_−33_(nt) makes one H-bond with S182 through its O6 atom; and the A_−31_(nt) makes one H-bond with M181. Moreover, M181 forms extensive van der Waals interactions with nucleotides at positions −31 and −30 of the template strand. The interactions are important, as alanine substitutions of R186 or S182 resulted in substantial loss of transcription activity (Figure 4C). Interestingly, the M181A mutant had increased transcription activity, but this came at the apparent expense of relaxing its sequence stringency for the positions −31 and −30 (Figure 4D), suggesting that M181 partially accounts for sequence specificity of the two positions.

The rest of the −35 element (−32, −30, and −29) makes no base-specific interactions. Previous crystal structure of *E. coli* σ^E^_4_/−35 dsDNA and a structural model of *S. coelicolor* σ^R^_4_/−35 dsDNA reported a local DNA shape readout (straight helix with a narrow minor groove) at this region^33, 42^. We observed a similar DNA conformation (Supplementary Figure 6B) in our crystal structure, suggesting a general mode of promoter recognition for the ECF σ factors. Such DNA structure might assist the binding of −35 dsDNA to the σ^H^_4_ surface perhaps by making favorable interactions through its phosphate backbones with polar residues of σ^H^_4_ including Y166, K167, T179, R183, H185, and R188 (Figure 3D and Supplementary Figure 6A). Consistently with this, losing any of these interactions causes a substantial loss of transcription activity (Figure 4C).

**Figure 4.**
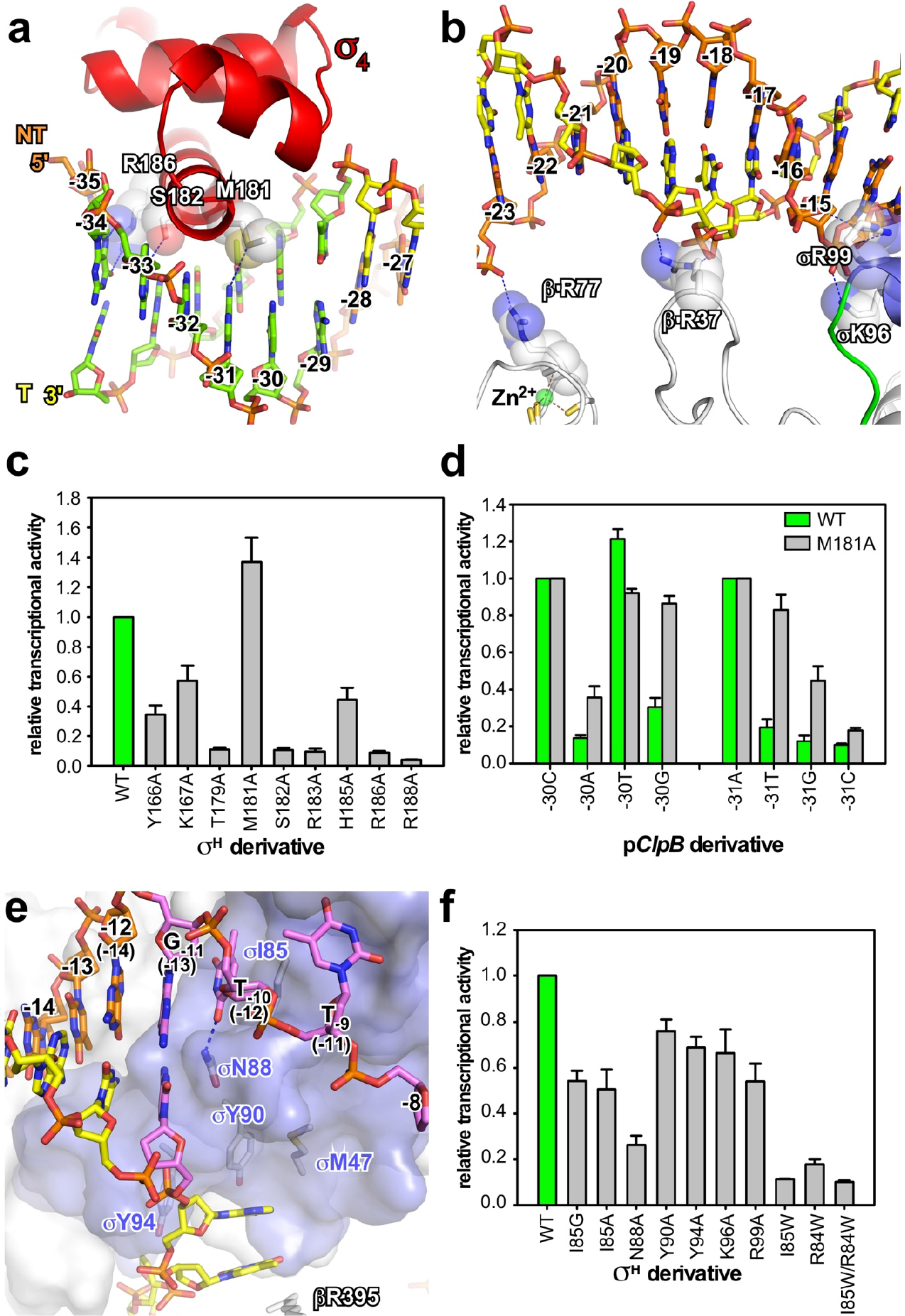
The interaction between upstream promoter DNA and RNAP in the *Mtb* σ^H^-RPo structure. **(A)** The interaction between σ^H^_4_ and the −35 element. σ^H^_4_, Red ribbon. The residues making base-specific interactions are presented as ribbon and half-transparent spheres; The C, O, N, and S atoms of the residues are colored in white, red, blue, and yellow respectively. The C, O, N, and P atoms of the −35 DNA are colored in green, red, blue, and orange, respectively. H-bond, blue dash. (B) The interaction between RNAP and −35/−10 spacer region of upstream dsDNA. The residues making polar interactions with DNA are shown and colored as above. The C atoms of the −35/−10 spacer DNA are colored in yellow (template DNA) or orange (nontemplate DNA), and the rest of atoms are colored as above. **(C)** The *in vitro* transcription activity of σ^H^ derivatives comprising alanine substitution on σ^H^_4_. **(D)** M181A changes the sequence specificity for positions −30 and −31 of the −35 element. **(E)** The promoter melting by σ^H^. The numbers in parenthesis correspond to positions in σ^A^-RPo. The −10 element is colored in purple. The residues involved in promoter melting are shown in stick and colored in white (C atoms), red (O atoms), and blue (N atoms). **(F)** The *in vitro* transcription activity of σ^H^_2_ derivatives with alanine or tryptophan substitutions. The experiments were repeated in triplicate, and the data are presented as mean ± S.E.M. Source data of (C), (D), and (F) are provided as a Source Data file.

### The interactions of σ^H^-RNAP with the −35/−10 spacer

In the crystal structure of *Mtb* σ^H^-RPo, σ^H^-RNAP contacts phosphate backbones of the spacer region between the −35 and the −10 elements at three positions (Figure 3D): 1) R77 of the RNAP-β′ zinc-binding domain (ZBD) contacts the nucleotide at position −23; 2) R37 of the RNAP-β′ zipper domain contacts the nucleotides at positions −20 and −19; and 3) K96 and R99 of σ^H^_2_ contact the nucleotide at position −14 (Figure 4B). These interactions probably stabilize the conformation of the upstream DNA duplex, likely promoting the engagement of the upstream duplex with σ^H^_4_ and σ^H^_2_ for subsequent promoter unwinding. Mutating K96 and R99 causes a mild loss of transcription activity, suggesting the importance of these interactions (Figure 4F).

### The promoter DNA unwinding function of σ^H^

The electron density map unambiguously shows that the T:A base pair at position −10 (corresponding to position −12 of promoters for σ^A^) is unwound, despite the fact that the −10 nucleotides in the synthetic nucleotide scaffold were designed to be complimentary (Figure 3D and 4E). This observation strongly suggests that σ^H^_2_ unwinds promoter DNA at the −11/−10 junction (corresponding to (−13)/(−12) of promoters targeted by σ^A^; Figure 4E and Supplementary Figure 6C-D). This clearly confirms the hypothesis that the ECF σ factors unwind promoter DNA starting from a distinct position as compared to σ^A 32, 34^. In the structure, N88 blocks the pathway of upstream dsDNA and serves as a wedge to disrupt the stacking of base pairs at the positions −11 and −10. The base pair at position −10 is subsequently forced open by σN88 via a competitive H-bond between the Watson-Crick atom of the T_−10_(nt) and N88 (Figure 4E). Two unwound nucleotides on the nontemplate strand DNA (T_−10_(nt) and T_−9_(nt)) are stabilized by two adjacent pockets of σ^H^_2_; these pockets are where the sequence identities are “read” (Supplementary Figure 6F). Moreover, two unwound nucleotides on the template strand DNA (A_−10_(t) and T_−9_(t)) are also trapped in a cleft created by the RNAP-β lobe and σ^H^_2_. Specifically, the base moieties of A_−10_(t) and T_−9_(t) form a stack with βR395, σY90, and σY94, and the phosphate moieties are stabilized by βK428, βN419, and σQ98. The functional importance of these residues for promoter DNA unwinding was underscored by our finding that their substitution with alanine resulted in defects in transcription (I85G, I85A, and N88A; Figure 4F).

Structure superimposition between σ^H^-RPo and σ^A^-RPo shows that the promoter DNA position at which unwinding is initiated differs between σ^A^ and σ^H^ by one base pair; σ^H^ unwinds promoter DNA at a position 1 bp upstream of the position at which σ^A^ unwinds its promoter DNA (the (−12)/(−11) junction for σ^A^; the −11/−10 junction for σ^H^ corresponding to the (−13)/(−12) junction for σ^A^; Supplementary Figure 6C-E). A tryptophan dyad (W433/W434 in *E. coli*; or W256/W257 in *T. aquaticus*) is essential for promoter unwinding at the (−12)/(−11) junction by σ^A 22, 23, 43, 44^, but the residues at the corresponding positions of σ^H^ (R84/I85) are not conserved (Supplementary Figure 3). Mutating R84 and I85 in σ^H^ to tryptophan (I85W, R84W, or I85W/R84W) resulted in substantial loss of transcription activity, confirming that σ^H^ opens promoter through a different mechanism than σ^A^ and supporting the mechanism proposed for *E. coli* σ^E^ (Figure 4F) ^34^.

### The interactions of σ^H^-RNAP with the −10 element

σ^H^-regulated promoters contain a ‘G_−11_T_−10_T_−9_’ consensus sequence at the −10 element (Supplementary Figure 5A) ^38, 40, 41^. Alteration of the DNA sequence at any of these positions resulted in complete loss of promoter activity (Supplementary Figure 5B), helping explain the reported finding from previous bioinformatic studies that the −10 element is the most conserved region among ECF σ factors ^10, 31^. Our crystal structure shows that the base moiety of C_−11_(t) of the −11 G:C pair makes one H-bond with N92 and extensive Van der Waal interactions with I91, alanine substitution of N92 or I91 resulted in modest or substantial decrease of transcription, respectively (Figure 5A and 5F), providing a structural explanation for sequence recognition at this position. Our crystal structure further reveals that σ^H^ recognizes the nucleotides of the next two positions via two protein pockets on the surface of σ^H^_2_ (Figure 5A).

**Figure 5.**
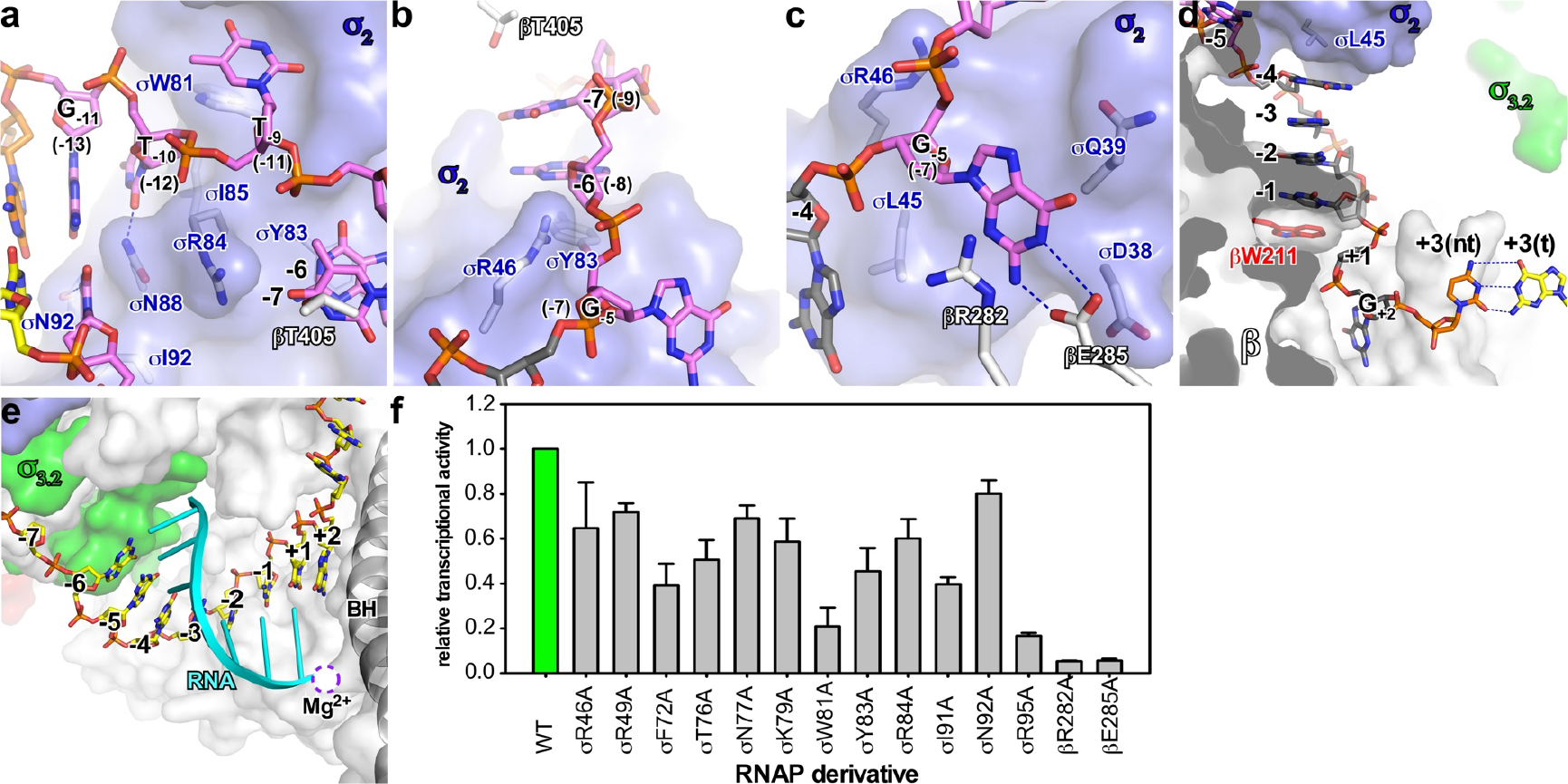
The interaction between transcription bubble and RNAP in the *Mtb* σ^H^-RPo structure. **(A)** Recognition of the ‘G_−11_T_−10_T_−9_’ sequence in the −10 element by σ^H^_2_. **(B)** Stacking of −7(nt) and −6(nt) nucleotides by RNAP-βsubunit and σ^H^_2_. **(C)** The recognition of G_−5_(nt) by RNAP-βsubunit and σ^H^_2_. **(D)** The interaction between RNAP and CRE element. Colors are as above. **(E)** The interaction between RNAP and DNA/RNA hybrid. **(F)** The *in vitro* transcription activity of RNAP derivatives comprising alanine substitutions of DNA-contacting residues on RNAP-βsubunit and σ^H^. The experiments were repeated in triplicate, and the data are presented as mean ± S.E.M. Source data of (F) are provided as a Source Data file.

T_−10_(nt) is accommodated by a shallow protein pocket on σ^H^_2_ wherein I85 forms a stack with the base of T_−10_(nt) at the pocket bottom and W81 supports the sugar moiety of T_−10_(nt) on one side of the pocket (Figure 5A). N88 on the other side of the pocket makes a H-bond with the base moiety of T_−10_(nt), likely contributing to the sequence specificity known to occur for this position. Alanine substitution of I85 or W81 causes severe defects in transcription activity (Figure 4F and 5F), emphasizing their importance. Sequence alignment of the ten *Mtb* ECF σ factors shows that the I85 and W81 are highly conserved (Supplementary Figure 3), suggesting the −10(nt) pocket probably exists on other ECF σ factors. Alanine substitution of N88 causes defects in transcription activity (Figure 4F), and the sequence alignment shows that N88 is the most frequent residue at this position. However, other polar residues occur at this position (*e.g.*, a histidine for σ^C^ and σ^D^, and an arginine for σ^I^, σ^J^, and σ^K^) (Supplementary Figure 3), suggesting that this position may help determine sequence specificity for position −10 of the promoter DNA.

T_−9_(nt) is flipped out and inserted into a protein pocket formed by the “specificity loop” of σ^H^_2_ ^34^. The thymine base of T_−9_(nt) stacked on top of W81 makes one H-bond with the main chain atom of σ^H^ residue R73 and two H-bonds with the side chains atoms of σ residues T76 and N77 on the specificity loop (Figure 3D and Supplementary Figure 6F). F72 and R73 contact the thymine base via van der Waals interactions. Mutating T_−9_(nt) completely abolished promoter activity (Supplementary Figure 5B), and alanine substitution of σ^H^ residues contacting T_−9_(nt) (F72A, T76A, N77A, and W81A) severely decreased transcription activity from the consensus promoter (Figure 5F), verifying the requirement for the T_−9_(nt)/σ^H^_2_ interaction for transcription. The fact that both the primary and the ECF σ factors use the specificity loop to read the sequence identity (position −9 of σ^H^-regulated promoters corresponding to position (−11) of σ^A^-regulated promoters; Supplementary Figure 6F-H) implies the central importance of this position in promoter DNA ^3, 24, 34^. Outside of these crucial positions, σ^H^_2_ forms fewer interactions with nontemplate nucleotides (positions −8, −7, and −6). In the structure, the base moiety of nucleotide at position −8 is disordered (no electron density) and the base moieties of −7 and −6 nucleotides are sandwiched between residue T405 of the RNAP-β subunit and residue Y83 of σ^H^_2_ (Figure 3D, 5B, and Supplementary Figure 6I).

A surprising finding in the σ^H^-RPo crystal structure is that σ^H^_2_ flips the guanine base of G_−5_(nt) (corresponding to the (−7) position of σ^A^-regulated promoters) and inserts into a shallow pocket created by σ^H^_2_ and the RNAP-β gate loop (Figure 5C). In this pocket, G_−5_(nt) forms a stacking interaction with R282, two H-bonds with E285 on the RNAP-β gate loop, and has van der Waals interactions with D38 and Q39 of σ^H^_2_. Mutations of the RNAP-β gate loop (βR282A or βE285A) cause severely-reduced transcription activity (Figure 5F). As the RNAP-β gate loop makes base-specific contacts to G_−5_(nt), we tested whether this position exhibits sequence preference. *In vitro* transcription assays showed that promoters with C or G at this position have much higher transcription activity compared with T or A (Supplementary Figure 5B), suggesting a sequence preference of C~G>T~A at such position. Our results therefore show that in σ^H^-regulated promoters, the consensus sequence of the −10 elements is extended to ‘G_−11_T_−10_T_−9_N_−8_N_−7_N_−6_S_−5_’. It is worth noting that σ^A^ also accommodates the T_(−7)_(nt) nucleotide in a protein pocket ^3, 24^; however the protein pocket is mainly formed by residues from σ^A^_1.2_ and σ^A^_2_^24^, and the structural features that determine sequence specificity are located on σ^A^ (Supplementary Figure 6I-K).

### Interactions of σ^H^-RNAP with the CRE

The guanine base of G_+2_(nt) is inserted into the ‘G’ pocket in σ^H^-RPo (Figure 5D) and makes essentially the same interaction with residues in the ‘G’ pocket as the G_(+2)_(nt) does in the σ^A^-RPo complex (Figure 3D, 5D, and Supplementary Figure 7D-F)^24^. However, in contrast to the σ^A^-RPo, in which the nontemplate T_(+1)_(nt) forms a stacking interaction with βW211, the nontemplate T_+1_(nt) in σ^H^-RPo was pushed out from the base-stacking. Instead, the nucleotide immediately upstream of −1 makes the stacking interaction with βW211 in σ^H^-RPo (Figure 5D and Supplementary Figure 7A-C). The base moieties of the remaining nucleotides (−4 to −1) between the CRE and −10 elements are stacked in a row between L45 on σ^H^_2_ and W211 on the RNAP-β subunit (Figure 5D and Supplementary Figure 7A).

### Interactions of the template ssDNA

In the σ^H^-RPo structure, σ^H^ and the RNAP-β subunit form an “T-ssDNA entry channel” that guides the template ssDNA into the RNAP active center cleft (Supplementary Figure 4A). Along the channel, σ^H^ and RNAP form extensive interactions with the template ssDNA (σM47, σR49, σY90, σY94, σQ98, σY04, βR395, βN419, βP422, βK428, and β′R334; Figure 3D). Compared to σ^A^, which forms extensive interactions between the σ^A^_3.2_ finger with the ssDNA template nucleotides the active center cleft ^24^, σ^H^ forms fewer interactions (Figure 3D and Supplementary Figure 7G-I). The 5’ terminus of the 7 nt RNA is positioned very closely to the tip of σ^H^_3.2_; extending the RNA by even one additional nucleotide would likely result in steric hindrance (Figure 5E and Supplementary Figure 7G). Since σ^H^_3.2_ occupies the RNA exit channel and must be displaced by the nascent RNA chain, such hindrance may be the trigger for the release of σ^H^ and promoter DNA during promoter escape.

## Discussion

The structural basis of transcription initiation by the primary σ factor has been studied extensively, but little is known about how the ECF σ factors—the largest and most diverse group of σ^70^ family factors—initiate transcription. In this study, we present high-resolution crystal structures of *M. tuberculosis* σ^H^-RNAP holoenzyme and σ^H^-RPo complexes along with comprehensive mutational analyses. Our study demonstrates the structural basis for RNAP holoenzyme formation and transcription initiation by the ECF σ factors.

Our structures show that σ^H^ binds to RNAP in a similar way to σ^A^, in which σ2 and σ_4_ stay on the surface of RNAP, and σ_3.2_ inserts into the active center. The interactions of σ^H^_2_ and σ^H^_4_ with RNAP were thus anticipated and supported a structure model of *E. coli* σ^E^-RNAP holoenzyme^45^; as the residues contacting the βFTH and β′CC domains are conserved between the ECF and primary σ factors. However, the interactions of σ^H^_3.2_ with RNAP are unexpected, showing similarity in neither sequence nor secondary structure between the σ_3.2_ regions of σ^ECF^ and σ^A^ (Supplementary Figure 3). *In vitro* transcription experiments show that σ^H^_3.2_ is essential for the transcription activity of σ^H^-RNAP; removing or replacing the linker with an unrelated sequence completely abolished its transcription activity (Figure 2D). It is worth noting that the B-reader loop of TFIIB reaches into the active site cleft of yeast pol II in a similar way to σ_3.2_ ^46^. Considering that this general mode of interaction appears to be conserved between prokaryotic RNAP and eukaryotic Pol II, it is reasonable to propose that the σ_2_/σ_4_ linker of other bacterial ECF σ factors very likely also inserts into the active site cleft of RNAP. Our transcription assays show that chimeric ECF σ factors with swapped σ_3.2_ domains retain function in transcription, albeit with reduced activity (Figure 2D and Supplementary Figure 4E), supporting this idea. An intriguing question to be answered is how RNAP uses the same channels to accommodate different σ_3.2_ domains.

σ^A^-RPo crystal structures show that σ^A^_3.2_ contacts nucleotides at the template strand of ssDNA and pre-organizes it into an A-form helical conformation in a manner compatible with pairing of initial NTPs ^24, 26^. These interactions provide explanations for the effects of σ^A^_3.2_ on *de novo* RNA synthesis ^26, 47, 48^. The structure predicts that the σ^A^_3.2_ finger has to be displaced by an RNA molecule of > 4nt length and that the σ^A^_3.2_ loop in the RNA exit channel has to be cleared during the promoter escape process. The interactions observed in the σ^A^-RPo structure underscore the key role of the σ^A^_3.2_ on abortive production, pausing, and promoter escape in transcription initiation ^28, 29, 49, 50^. In our crystal structures of σ^H^-RPo, we show that σ^H^_3.2_ guides the template ssDNA into the active site cleft and forms interactions with template ssDNA (Figure 3D and 5E). We propose that σ^H^_3.2_ probably functions similarly to σ^A^_3.2_ during transcription initiation: by stabilizing the template ssDNA and facilitating binding of initial NTPs. The crystals structure of σ^H^-RPo also indicates that σ^H^_3.2_ should collide with RNA molecules > 7 nt in length and that the σ^H^_3.2_ domain must dissociate from the RNAP RNA exit channel during promoter escape (Figure 5E), raising the possibility that the σ_3.2_ of the ECF factors functions like σ^A^_3.2_ during abortive production, pausing, and promoter escape in transcription initiation.

Our structure of σ^H^-RPo suggests that substantial differences exist between how individual domains σ^H^-RNAP and σ^A^-RNAP interact with their cognate promoter DNA. Both σ^H^_4_ and σ^A^_4_ use the same α-helix to bind the −35 element, but the positions of DNA on the α-helix differ by one α-helical turn, resulting a ~ 4 Å difference in the position of the −35 element on the σ_4_ surface (Supplementary Figure 6B). A previous crystal structure of the *E. coli* σ^E^_4_/−35 element binary complex is superimposable on our σ^H^-RPo (Supplementary Figure 6B)^33^, suggesting that the distinct mode of interaction that we observed with the −35 element is likely used by other ECF σ factors.

σ^H^-RNAP **“** reads” the sequence of the −10 element differently than does σ^A^-RNAP. In our crystal structure of σ^H^-RPo, we discovered that base moieties of three nucleotides—T_−10_(nt), T_−9_(nt), and G_−5_(nt) (corresponding to the positions (−12), (−11) and (−7) of σ^A^-regulated promoters)—were flipped out and inserted into three respective protein pockets on σ^H^ (Figure 3D and 5A-C), in contrast to the two protein pockets known for base moieties of A_(−11)_(nt) and T_(−7)_(nt) on σ^A 3, 24^. This extra pocket for T_−10_(nt) on σ^H^ was also suggested in a previous structure of a *E. coli* σ^E^_2_/−10 element binary complex (Supplementary Figure 6G)^34^. Sequence alignment of multiple ECF σ factors and σ^A^ revealed that residues forming the T_−10_(nt) pocket are generally conserved between ECF σ factors but are distinct from σ^A^, suggesting that other ECF σ factors likely also recognize the nucleotide at this position using similar protein pockets (Supplementary Figure 3).

σ^H^-RNAP uses different protein regions to accommodate the flipped guanine base of G_−5_(nt) (corresponding to position (−7) of σ^A^-regulated promoters) than does σ^A^-RNAP for T_(−7)_(nt) (Supplementary Figure 6I-K). The guanine base of G-_5_(nt) is sandwiched between σ^H^_2_ and the RNAP-β gate loop, while the thymine base of T_(−7)_(nt) resides in a pocket on σ^A^_1.2_^3, 24^. Our mutation study of the G-_5_(nt) pocket residues demonstrated that the RNAP-β gate loop functions to recognize this particular nucleotide (Figure 5F), thus raising the possibility that other σ^ECF^-RNAP holoenzymes may also bind and read a nucleotide in the nontemplate ssDNA in a manner analogous to σ^H^-RNAP.

σ^H^-RNAP also engages the −10 element differently than does σ^A^-RNAP. We found that the protein pockets for T_−9_(nt) and G_−5_(nt) on σ^H^-RNAP do not exist in the absence of promoter DNA (Figure 6A-B). In the crystal structure of σ^H^-RNAP: the specificity loop which recognizes the T_−9_(nt) is disordered; and the RNAP-β gate loop is too far away from the σ^H^_2_ to form the G_−5_(nt) pocket (Figure 6A-B). Such conformational differences support an “induced fit” model of interaction between σ^H^-RNAP and nontemplate ssDNA, in contrast to the accepted “lock-and-key model” for the interaction between σ^A^-RNAP and nontemplate ssDNA (Figure 6C-D) ^3, 24, 51^.

**Figure 6.**
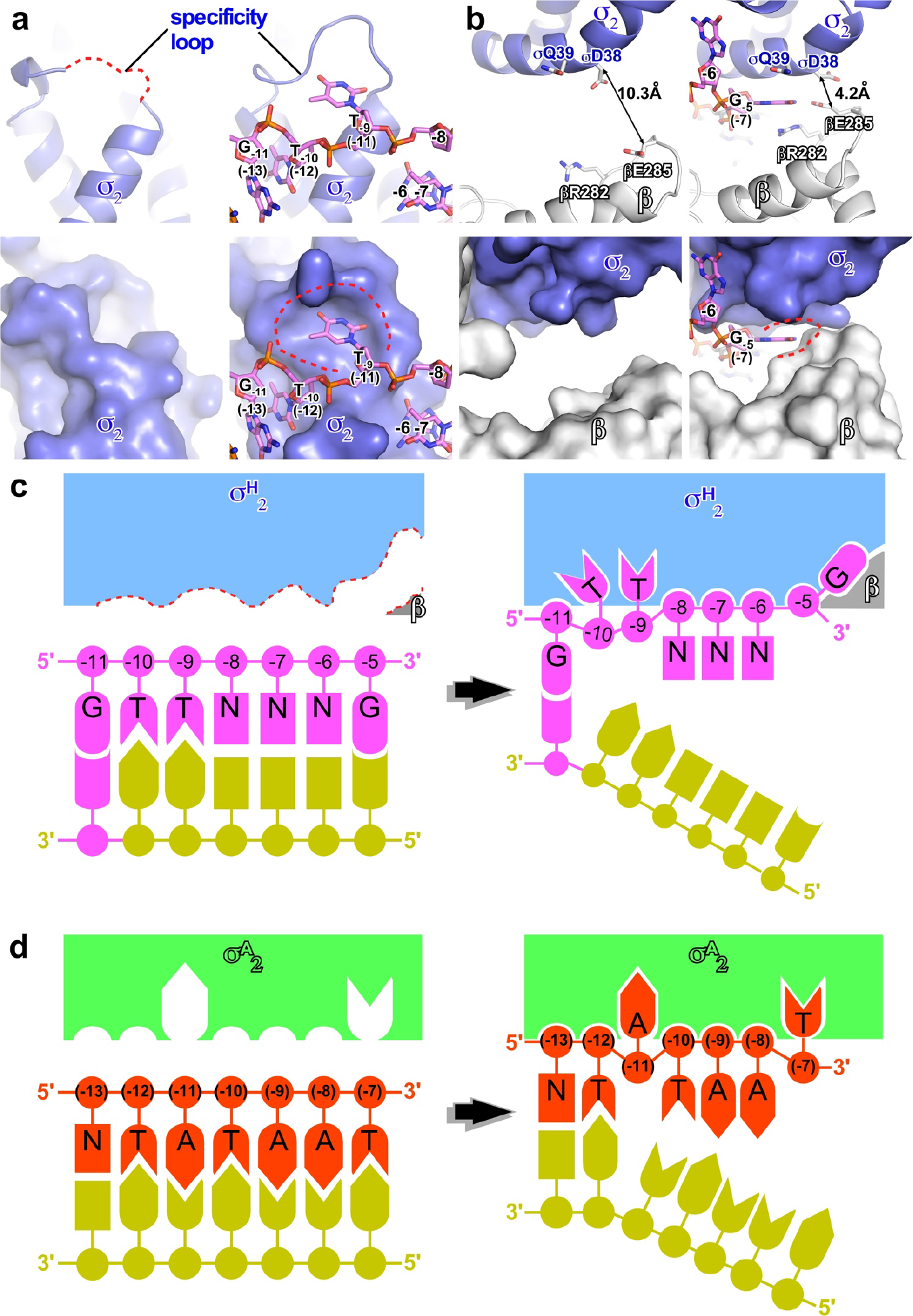
The induced-fit mechanism of promoter recognition by *Mtb* σ^H^-RNAP. **(A)** The T_−9_(nt) pocket does not exist in σ^H^-RNAP holoenzyme (left) but is induced by DNA binding in σ^H^-RPo (right) structures. **(B)** The G_−5_(nt) pocket does not exist in σ^H^-RNAP holoenzyme (left) but is induced by DNA binding in σ^H^-RPo (right) structures. The pockets are presented in cartoon (top) and surface (bottom). **(C)** The schematic of induce-fit model of promoter recognition by σ^H^-RNAP. **(D)** The schematic of lock-and-key model of promoter recognition by σ^A^-RNAP.

σ^H^-RNAP recognizes the G_+2_(nt) of CRE in a same way as does σ^A^-RNAP (Supplementary Figure 7D-F). As the residues that form the ‘G’ pocket are solely from the RNAP core enzyme, it is possible that other ECF σ-RNAP holoenzymes are probably able to read the sequence identity of nucleotide at position +2 of the promoter DNA. However, whether the sequence content at this position affects other events (transcription start site selection, slippage synthesis, *etc.*) during transcription initiation by ECF σ-RNAP as σ^A^-RNAP remains to be determined ^52^.

Our crystal structures suggest that σ^H^ employs a distinct mechanism to unwind promoter DNA compared to σ^A^ (Supplementary Figure 6C-E): 1) σ^H^ and σ^A^ use residues with positions that differ by one α-helical turn on the σ_2.3_ α-helix (N88 for *Mtb* σ^H^ *vs.* W433/W434 for *Ec* σ^A^, or W256/W257 for *Taq* σ^A^) to unwind promoter DNA; 2) σ^H^ and σ^A^ unwind promoter DNA at positions differing by one base pair ((−13)/(−12) junction for σ^H^ *vs.* −(12)/(−11) junction for σ^A^); and 3) σ^H^ traps and reads two unwound nucleotides (T_(−12)_(nt) and T_(−11)_(nt) immediately after the unwinding points), whereas σ^A^ only traps and reads one unwound nucleotide (A_(−11)_(nt)). Although it is unclear whether trapping of the flipped nucleotides initiates or facilitates the event of promoter unwinding, such interactions play crucial roles during RPo formation.

Campagne *et al.* 2014 recently identified a similar protein pocket on *E. coli* σ^E^_2_ for the T_(−12)_(nt) in a crystal structure of *E. coli* σ^E^ bound to the −10 element ssDNA, and predicted that *E. coli* σ^E^ unwinds promoter dsDNA at the (−13)/(−12) junction^34^. Our crystal structure of σ^H^-RPo clearly confirms the unwinding position proposed in the Campagne *et al.* 2014 study. Sequence alignment of multiple ECF σ factors and σ^A^ revealed that most of the ECF σ factors do not contain the tryptophan dyad of σ^A^ at corresponding positions, but instead share a conserved (−12) pocket (Supplementary Figure 3). Therefore, it is possible that the ECF σ factors share the same unwinding mechanism as *Mtb* σ^H^ and *Ec* σ^E^.

Our mutation study of the σ^H^-regulated promoter showed that substitution of the consensus sequence at almost every position on the −35 element and −10 element abolished transcription activity (Supplementary Figure 5B). Moreover, extending or shortening the spacer of −35/−10 elements substantially reduced promoter activity (Figure 2B). These results confirmed previously reported observations that ECF σ factors require a consensus sequence at the −35/−10 elements as well as a rigid spacer on promoter DNA to efficiently initiate transcription ^8, 32^. Our structures and results from biochemical experiments provide explanations for the promoter stringency of σ^H^. We show that the interactions among σ^H^ and RNAP, the unwinding mechanism, and the induced-fit mode of promoter recognition work in concert to collectively confer the high specificity exhibited by σ^H^ and probably by other ECF σ factors as well.

We have shown that σ^H^ employs residues different from σ^A^ to unwind promoter DNA. The well-conserved tryptophan dyad of σ^A^ functions very efficiently for promoter unwinding; substitutions of the tryptophan dyad in σ^A^ resulted in severely reduced transcription activity ^43, 44^; and sequence variations at corresponding positions accounts for inferior DNA unwinding capacity of other alternative σ factors ^25^. Given that ECF σ factors lacks the tryptophan dyad at corresponding positions, we infer σ^H^ (and probably other ECF σ factors) unwinds promoter DNA less efficiently than does σ^A^. This putative sub-optimal unwinding efficiency could be compensated by employing a very high affinity consensus sequence of promoter DNA to facilitate its loading ^4, 8, 25^. Our proposed induced-fit mode of nontemplate ssDNA binding by σ^H^-RNAP at the position immediate downstream of unwinding--*i.e.* T_−9_(nt)--also require the consensus promoter sequence to induce formation of correct conformation of the ‘specificity loop’ (Figure 6C); RNAP is not able to efficiently propagate promoter unwinding downstream without firmly anchoring the ‘master’ nucleotide-- A_−11_(nt) for *E. coli* σ^70^ corresponding to T-_9_(nt) for *Mtb* σ^H^ --as demonstrated in the case of *E. coli* σ^70^-RNAP ^53, 54, 55^.

In conclusion, we demonstrate the structural basis of RNAP holoenzyme formation and transcription initiation by the ECF σ factors, thereby deepening our understanding the basic mechanisms of transcription initiation used by the largest and most diverse group of bacterial initiation factors. Our work will facilitate the rational design of orthogonal transcription units based on ECF σ factors and should help computational chemistry and other efforts to design selective antibacterial agents through the inhibition ECF σ factor mediated transcription initiation.

## Methods

### Plasmid construction

The plasmids used in this study is listed in Supplementary 1. For construction of the expression plasmid pTolo-EX5-*Mtb*σ^H^, the *M. tuberculosis* σ^H^ gene amplified from *M. tubercolusis* genomic DNA (see Supplementary Data 1 for primer information) was cloned into the pTolo-EX5 plasmid (Tolo Biotech.) using NcoI and XhoI restriction sites. The pTolo-EX5-*Mtb*σ^H^ derivatives bearing single or double mutations were generated through site-directed mutagenesis (Transgen biotech).

The pTolo-EX5-*Mtb*σ^H^ derivatives encoding chimeric σ^H^ were generated by replacing the DNA fragment encoding *Mtb* σ^H^_3.2_ (aa 96-144) with DNA fragments encoding *Ec* σ^A^ (aa 164-212; disordered acidic loop of the non-conserved region), *Mtb* σ^E^_3.2_ (aa 150-189), *Mtb* σ^L^_3.2_ (aa 78-122), or *Mtb* σ^M^_3.2_ (aa 98-137) in pTolo-EX5-*Mtb*σ^H^ (Tolo Biotech).

The pACYCduet-*Mtb*-*rpoA-rpoZ* plasmid was constructed by replacing *Mtb rpoD* with *Mtb rpoZ* in parent plasmid pACYC-duet-*Mtb*-*rpoA-sigA* plasmid using KpnI and NdeI (Supplementary Table 1). The pETduet-*Mtb*-*rpoB-rpoC* derivatives bearing single mutations were generated through site-directed mutagenesis (Transgen biotech; Supplementary Table 1 and Supplementary Data 1).

For construction of plasmids for *in vitro* transcription assays of *Mtb* σ^H^, the promoter region (−50 to +51) of *ClpB* gene amplified from *M. tuberculosis* genomic DNA was cloned into pEASY-Blunt simple vectors, resulting in pEASY-Blunt-p*ClpB* (Transgen biotech; Supplementary Table 1 and 2). The derivatives of pEASY-Blunt-p*ClpB* with varied −35/−10 spacer lengths were obtained by site-directed mutagenesis (Supplementary Figure 2J). The promoter region (−50 to +51) of *Rv2466c* gene amplified from *M. tuberculosis* genomic DNA was cloned into pEASY-Blunt simple vectors, resulting in pEASY-Blunt-p*Rv2466c* (Supplementary Table 1 and Supplementary Data 1; Supplementary Figure 2L).

The derivatives of pARTaq-N25-100-TR2 for *in vitro* transcription assays of *Mtb* σ^A^ with varied −35/−10 spacer lengths were obtained by site-directed mutagenesis (Supplementary Figure 2K).

### Protein preparation

For preparation of *M. tuberculosis* σ^H^, *E. coli* BL21(DE3) cells (NovoProtein) carrying pTolo-EX5-*Mtb*σ^H^ were cultured in LB at 37 ºC, and the expression of N-terminal sumo-tagged *Mtb* σ^H^ was induced at 18 ºC for 14h with 0.5 mM IPTG at OD_600_ of 0.8. Cells were harvested by centrifugation (8,000 g, 4 ºC), re-suspended in lysis buffer (20 mM Tris-HCl pH 8.0, 0.5 M NaCl, 5% (v/v) glycerol, 0.5 mM β-mercaptoethanol, protease inhibitor cocktail (bimake.cn)) and lysed using an Avestin EmulsiFlex-C3 cell disrupter (Avestin, Inc.). The lysate was centrifuged (16,000 g; 45 min, 4ºC) and the supernatant was loaded on to a 2 ml column packed with Ni-NTA agarose (SMART, Inc.). The protein was washed by lysis buffer containing 20 mM imidazole and eluted with lysis buffer containing 250 mM imidazole. The eluted fraction was digested by TEV protease and dialyzed overnight in dialysis buffer (20 mM Tris-HCl pH 8.0, 0.2 M NaCl, 1% (v/v) glycerol, 0.5 mM β-mercaptoethanol). The sample was loaded onto a second Ni-NTA column and the cleaved protein was retrieved from the flow-through fraction. The sample was diluted to the dialysis buffer with 0.05 M NaCl and further purified through a Heparin column (HiTrap Heparin HP 5ml column, GE healthcare Life Sciences) with buffer A (20 mM Tris-HCl pH8.0, 0.05 M NaCl, 1% (v/v) glycerol, 1 mM DTT) and buffer B (20 mM Tris-HCl pH 8.0, 1 M NaCl, 1% (v/v) glycerol, 1 mM DTT). Fractions containing *M. tuberculosis* σ^H^ was concentrated to 5 mg/ml and stored at −80 ºC. The *M. tuberculosis* σ^H^ derivatives were prepared by the same procedure.

For preparation of selenomethionines (SeMet)-labeled *M. tuberculosis* σ^H^, BL21 (DE3) strains carrying pTolo-EX5-*Mtb*σ^H^ were cultured in SelenoMet base medium supplemented with nutrient mix (Molecular Dimensions) at 37 ºC. The amino acids mixture containing Selemethionine were added into the culture at OD_600_ of 0.4 and the protein expression was induced with 0.5 mM IPTG at OD_600_ of 0.8 for 14h at 18 ºC. The SeMet-labeled *Mtb* σ^H^ was purified as described above.

The *M. tuberculosis* RNA polymerase core enzyme was expressed and purified from *E. coli* BL21(DE3) carrying pETDuet-*Mtb-rpoA-rpoZ* and pACYCDuet-*Mtb*-*rpoB-rpoC* as described^56^. The protein sample was concentrated to 5 mg/ml and stored at −80 ºC.

### Nucleic-acid scaffolds

Nucleic acid scaffolds for assembly of σ^H^-RPo* for crystallization of σ^H^-RNAP holoenzyme was prepared from synthetic oligos (nontemplate DNA: 5’-GTTGTGCTGGGCGTCACGGATGCA-3’; template DNA: 5’- TGCATCCGTGAGTCGGT-3’, Sangon Biotech, Supplementary Figure 1A) by an annealing procedure (95°C, 5 min followed by 2°C-step cooling to 25°C) in annealing buffer (5 mM Tris-HCl, pH 8.0, 200 mM NaCl, and 10 mM MgCl2).

Nucleic acid scaffolds for crystallization of σ^H^-RPo was prepared from synthetic oligos (nontemplate DNA: 5’- CGGAACAGTTGCGACTTAGACGTGGTTGTGGGAGCTGCTATACTCTCC -3’; template DNA: 5’- GGAGAGTATAGGTCGAGGGTGTACCACGTCTAAGTCGCAACTGTTCC -3’, Sangon Biotech; and RNA: 5’- CCCUCGA -3’, Genepharma; Figure 3C) by an annealing procedure (95°C, 5 min followed by 2°C-step cooling to 25°C) in annealing buffer (5 mM Tris-HCl, pH 8.0, 200 mM NaCl, and 10 mM MgCl_2_).

### *M. tuberculosis* σ^H^-RPo complex reconstitution

The *M. tuberculosis* σ^H^-RPo and σ^H^-RPo*was reconstituted from *M. tuberculosis* RNAP core enzyme, σ^H^ (or SeMet-σ^H^) and nucleic acid scaffolds. The RNAP core enzyme, σ^H^ and nucleic acid scaffolds were mixed at a 1:4:1.2 molar ratio and incubate at 4ºC overnight. The mixture was loaded on a HiLoad 16/60 Superdex S200 column (GE Healthcare, Inc.) equilibrated in 20 mM Tris-HCl pH 8.0, 0.1 M NaCl, 1%(v/v) glycerol, 1 mM DTT. Fractions containing *Mtb* σ^H^-RPo were collected, concentrated to 7.5 mg/ml, and stored at −80ºC.

### Structure determination *M. tuberculosis* σ^H^-RNAP holoenzyme

The structure of σ^H^-RNAP holoenzyme was obtained during an attempt for obtaining the σ^H^-RPo* with the fork transcription bubble DNA scaffold (no RNA oligo in the scaffold). The initial screen was performed by a sitting-drop vapor diffusion technique. Crystals grown from optimized reservoir solution A (1 μL 0.2 M NaAc, 0.1 M sodium citrate pH5.5, 10% PEG4000 mixed with 1μL 7.5 mg/mL protein complex) for 3 days at 22 ºC were harvested for X-ray diffraction data collection. Crystals were soaked in stepwise fashion to reservoir solution A containing 18%(v/v) (2R, 3R)-(−)-2,3-butanediol (Sigma-Aldrich) and cooled in liquid nitrogen. The crystals of σ^H^-RNAP derivative containing SeMet-labeled σ^H^ were obtained by analogous procedure.

Data were collected at Shanghai Synchrotron Radiation Facility (SSRF) beamlines 17U and 19U1, processed using HKL2000 ^57^. The structure was solved by molecular replacement with Phaser MR^58^ using the structure of *M. smegmatis* core enzyme in a *M. smegmatis* transcription initiation complex (PDB:5TW1) ^35^ [https://www.rcsb.org/structure/5TW1] as the search model. Only one molecule of RNAP core enzyme was found in one asymmetric unit. The electron density maps show clear signal for σ^H^. Cycles of iterative model building and refinement were performed in Coot ^59^ and Phenix^60^. Residues of σ^H^ were built into the model at the last stage of refinement. No density of nucleic acid was observed in all stages of refinements, suggesting that the nucleic acids dissociated during crystallization resulting in a crystal of σ^H^-RNAP holoenzyme. The final model of *Mtb* σ^H^-RNAP holoenzyme was refined to R_work_ and R_free_ of 0.218 and 0.258, respectively. Analogous procedures were used to refine the structures of σ^H^-RNAP holoenzyme with SeMet-labeled σ^H^.

### Structure determination *M. tuberculosis* σ^H^-RPo

The initial screen of σ^H^-RPo was performed by a sitting-drop vapor diffusion technique. Crystals grown from reservoir solution B (1 μL 2% Tacsimate pH 5.0, 0.1 M Sodium citrate pH 5.6, 16 % PEG3350 mixed with 1μL 7.5 mg/mL protein complex) for 15 days at 22ºC were harvested for X-ray diffraction data collection. Crystals were soaked in stepwise fashion to the reservoir solution B containing 18%(v/v) (2R, 3R)-(−)-2,3-butanediol (Sigma-Aldrich) and cooled in liquid nitrogen. Data were collected at Shanghai Synchrotron Radiation Facility (SSRF) beamlines 17U and 19U1, processed using HKL2000 ^57^. The structure was solved by molecular replacement with Phaser MR^58^ using the structure of *M. tuberculosis* σ^H^-RNAP as a search model. Only one molecule of σ^H^-RNAP was found in one asymmetric unit. The electron density map showed clear signals for nucleotides in transcription bubble and downstream DNA duplex after initial rigid-body refinement, and clear signals for nucleotides in upstream DNA duplex after iterative cycles of model building and refinements in Coot ^59^ and Phenix^60^. The nucleotides were built into the model at the last stage, and the final model of *Mtb* σ^H^-RPo was refined to R_work_ and R_free_ of 0.220 and 0.255, respectively.

### *In vitro* transcription assay

Transcription assays with *M. tuberculosis* RNAP σ^H^-holoenzyme were performed as follows: Reaction mixtures contained (20 μL): 80 nM *M. tuberculosis* RNAP core enzyme, 1 μM *M. tuberculosis* σ^H^, 40 mM Tris-HCl, pH 7.9, 75 mM KCl, 5 mM MgCl_2_, 2.5 mM DTT, and 12.5% glycerol. Reaction mixtures were incubated 10 min at 37°C, and then supplemented with 2 μl promoter DNA (1 μM; amplified from pEASY-Blunt-p*ClpB;* Supplementary Data 1), and further incubated 10 min at 37°C. The reaction were initiated by adding 0.7 μl NTP mixture (3 mM [α-^32^P]UTP (0.04 Bq/fmol), 3 mM ATP, 3 mM GTP, and 3 mM CTP), and RNA synthesis was allowed to proceed for 10 min at 37°C. Reactions were terminated by adding 8 μl loading buffer (10 mM EDTA, 0.02% bromophenol blue, 0.02% xylene cyanol, and 98% formamide), boiled for 2 min, and stored in ice for 5 min. Reaction mixtures were applied to 15% urea-polyacrylamide slab gels (19:1 acrylamide/bisacrylamide), electrophoresed in 90 mM Tris-borate (pH 8.0) and 0.2 mM EDTA, and analyzed by storage-phosphor scanning (Typhoon; GE Healthcare, Inc.).

Transcription assays with *M. tuberculosis* RNAP σ^A^-holoenzyme were performed essentially as above except that σ^A^ instead of σ^H^ were added and N25 promoter DNA were used (amplified from pARTaq-N25-100-TR2; Supplementary Table 1 and Supplementary Data 1).

Transcription assays using with *M. tuberculosis* RNAP σ^H^-holoenzyme and p*Rv2466c* promoters were also performed essentially as above with subtle modifications. The reaction mixtures (20 μL) containing 160 nM *M. tuberculosis* RNAP core enzyme, 1 μM *M. tuberculosis* σ^H^, 40 mM Tris-HCl, pH 7.9, 75 mM KCl, 5 mM MgCl_2_, 2.5 mM DTT, and 12.5% glycerol were incubated 10 min at 37 °C, and then supplemented with 2 μl promoter DNA (1 μM, amplified from pEASY-p*Rv2466*c; Supplementary Table 1 and Supplementary Data 1), and further incubated for 10 min at 37°C. The reactions were initiated by adding 4 μl NTP mixture (0.1 mM ATP, 0.1 mM GTP, 0.1 mM CTP, and 7 μM [α-^32^P]UTP (5.6 Bq/fmol) and were allowed to proceed for 20 min at 37 °C. The reactions were terminated and the transcripts were separated and visualized as above.

### Quantification and statistical analysis

All biochemical assays were performed at least 3 times independently. Data were analyzed with SigmaPlot 10.0 (Systat Software Inc.).

### Data availability

The accession number for the coordinates and structure factors for *M. tuberculosis* σ^H^-RNAP and σ^H^-RPo in this paper are PDB: 5ZX3 and 5ZX2, respectively. The source data underlying Figures 2B, 2D, 4C, 4D, 4F, 5F, and Supplementary Figures 2I, 4E, and 5B are provided as a Source Data file. Other data from the corresponding author upon reasonable request.

## Acknowledgements

Work was supported by National Natural Science Foundation of China (31670067 and 31822001), the Chinese Thousand Talents Program to Y.Z., the Strategic Priority Research Program of the Chinese Academy of Sciences (XDB29020000), and the Leading Science Key Research Program of the Chinese Academy of Sciences (QYZDB-SSW-SMC005). We thank Prof. Richard Ebright for generous gifts of pACYC-duet-*Mtb*-*rpoA*-*rpoD*, pETDuet-*Mtb-rpoB-rpoC*, and pARTaq-N25-100-TR2 constructs, Prof. Xiaoming Zhang for generous gift of *M. tuberculosis* genomic DNA, and Tolo Biotechnology for generous gift of pTolo-EX vectors. We thank the staff at beamline BL18U1/BL19U1 of National Center for Protein Science Shanghai (NCPSS), and at beamline BL17U1 of Shanghai Synchrotron Radiation Facility for assistance during data collection.

## Author contributions

L. L. solved the structures. C. F. performed *in vitro* transcription assays. N. Z. assisted in structure determination. T. W. prepared pACYCduet-*Mtb*-*rpo*A-*rpoZ* and purified proteins. Y. Z. designed experiments, analyzed data, and wrote the manuscript.

## Supplementary Information

Structural basis for transcription initiation by bacterial ECF σ factors

Lingting Li, Chengli Fang *et al.*

**Supplementary Table 1.** Plasmids used in this study.

| REAGENT | SOURCE | REAGENT | SOURCE |
| --- | --- | --- | --- |
| pTolo-EX5 | Tolo Biotech | pTolo-EX5-*Mtb*σ<sup>H</sup>-S182A | This study |
| pTolo-EX5-*Mtb*σ<sup>H</sup> | This study | pTolo-EX5-*Mtb*σ<sup>H</sup>-R183A | This study |
| pTolo-EX5-*Mtb*σ<sup>H</sup>-R46A | This study | pTolo-EX5-*Mtb*σ<sup>H</sup>-H185A | This study |
| pTolo-EX5-*Mtb*σ<sup>H</sup>-R49A | This study | pTolo-EX5-*Mtb*σ<sup>H</sup>-R186A | This study |
| pTolo-EX5-*Mtb*σ<sup>H</sup>-F72A | This study | pTolo-EX5-*Mtb*σ<sup>H</sup>-R188A | This study |
| pTolo-EX5-*Mtb*σ<sup>H</sup>-T76A | This study | pTolo-EX5-*Mtb*σ<sup>H</sup><sub>2</sub>(1-95) | This study |
| pTolo-EX5-*Mtb*σ<sup>H</sup>-N77A | This study | pTolo-EX5-*Mtb*σ<sup>H</sup>-I85W/R84W | This study |
| pTolo-EX5-*Mtb*σ<sup>H</sup>-K79A | This study | pTolo-EX5-*Mtb*σ<sup>H</sup><sub>4</sub>(145-216) | This study |
| pTolo-EX5-*Mtb*σ<sup>H</sup>-W81A | This study | pTolo-EX5-*Mtb*σ<sup>H</sup><sub>2</sub>-DL-σ<sup>H</sup><sub>4</sub> | This study |
| pTolo-EX5-*Mtb*σ<sup>H</sup>-Y83A | This study | pTolo-EX5-*Mtb*σ<sup>H</sup><sub>2</sub>-σ<sup>E</sup><sub>3.2</sub>-σ<sup>H</sup><sub>4</sub> | This study |
| pTolo-EX5-*Mtb*σ<sup>H</sup>-R84A | This study | pTolo-EX5-*Mtb*σ<sup>H</sup><sub>2</sub>-σ<sup>L</sup><sub>3.2</sub>-σ<sup>H</sup><sub>4</sub> | This study |
| pTolo-EX5-*Mtb*σ<sup>H</sup>-R84W | This study | pTolo-EX5-*Mtb*σ<sup>H</sup><sub>2</sub>-σ<sup>M</sup><sub>3.2</sub>-σ<sup>H</sup><sub>4</sub> | This study |
| pTolo-EX5-*Mtb*σ<sup>H</sup>-I85A | This study | pACYCduet-*Mtb-rpoA-rpoZ* | This study |
| pTolo-EX5-*Mtb*σ<sup>H</sup>-I85G | This study | pETduet-*Mtb-rpoB*(R282A)- | This study |
| pTolo-EX5-*Mtb*σ<sup>H</sup>-I85W | This study | pETduet-*Mtb-rpoB*(E285A)- | This study |
| pTolo-EX5-*Mtb*σ<sup>H</sup>-N88A | This study | pEASY-Blunt-p*ClpB*-spacer15 | This study |
| pTolo-EX5-*Mtb*σ<sup>H</sup>-I91A | This study | pEASY-Blunt-p*ClpB*-spacer16 | This study |
| pTolo-EX5-*Mtb*σ<sup>H</sup>-N92A | This study | pEASY-Blunt-p*ClpB*-spacer17 | This study |
| pTolo-EX5-*Mtb*σ<sup>H</sup>-K96A | This study | pEASY-Blunt-p*ClpB*-spacer18 | This study |
| pTolo-EX5-*Mtb*σ<sup>H</sup>-R99A | This study | pEASY-Blunt-p*ClpB*-spacer19 | This study |
| pTolo-EX5-*Mtb*σ<sup>H</sup>-Y166A | This study | pARTaq-N25-100-TR2- | This study |
| pTolo-EX5-*Mtb*σ<sup>H</sup>-K167A | This study | pARTaq-N25-100-TR2- | This study |
| pTolo-EX5-*Mtb*σ<sup>H</sup>-T179A | This study | pARTaq-N25-100-TR2- | This study |
| pTolo-EX5-*Mtb*σ<sup>H</sup>-M181A | This study | pARTaq-N25-100-TR2- | This study |
| pEASY-Blunt-p*ClpB*(-5A) | This study | pEASY-Blunt-p*Rv2466c* | This study |
| pEASY-Blunt-p*ClpB*(-5T) | This study | pEASY-Blunt-p*ClpB*(-30A) | This study |
| pEASY-Blunt-p*ClpB*(-5G) | This study | pEASY-Blunt-p*ClpB*(-30T) | This study |
| pEASY-Blunt-p*ClpB*(-9A) | This study | pEASY-Blunt-p*ClpB*(-30G) | This study |
| pEASY-Blunt-p*ClpB*(-9G) | This study | pEASY-Blunt-p*ClpB*(-31T) | This study |
| pEASY-Blunt-p*ClpB*(-9C) | This study | pEASY-Blunt-p*ClpB*(-31G) | This study |
| pEASY-Blunt-p*ClpB*(-10A) | This study | pEASY-Blunt-p*ClpB*(-31C) | This study |
| pEASY-Blunt-p*ClpB*(-10G) | This study | pEASY-Blunt-p*ClpB*(-32G) | This study |
| pEASY-Blunt-p*ClpB*(-10C) | This study | pEASY-Blunt-p*ClpB*(-32C) | This study |
| pEASY-Blunt-p*ClpB*(-11A) | This study | pEASY-Blunt-p*ClpB*(-32A) | This study |
| pEASY-Blunt-p*ClpB*(-11T) | This study | pEASY-Blunt-p*ClpB*(-33A) | This study |
| pEASY-Blunt-p*ClpB*(-11C) | This study | pEASY-Blunt-p*ClpB*(-33T) | This study |
| pEASY-Blunt-p*ClpB*(-29G) | This study | pEASY-Blunt-p*ClpB*(-33C) | This study |
| pEASY-Blunt-p*ClpB*(-29T) | This study | pEASY-Blunt-p*ClpB*(-34A) | This study |
| pEASY-Blunt-p*ClpB*(-29C) | This study | pEASY-Blunt-p*ClpB*(-34T) | This study |
|  |  | pEASY-Blunt-p*ClpB*(-34C) | This study |

**Supplementary Figure 1.**
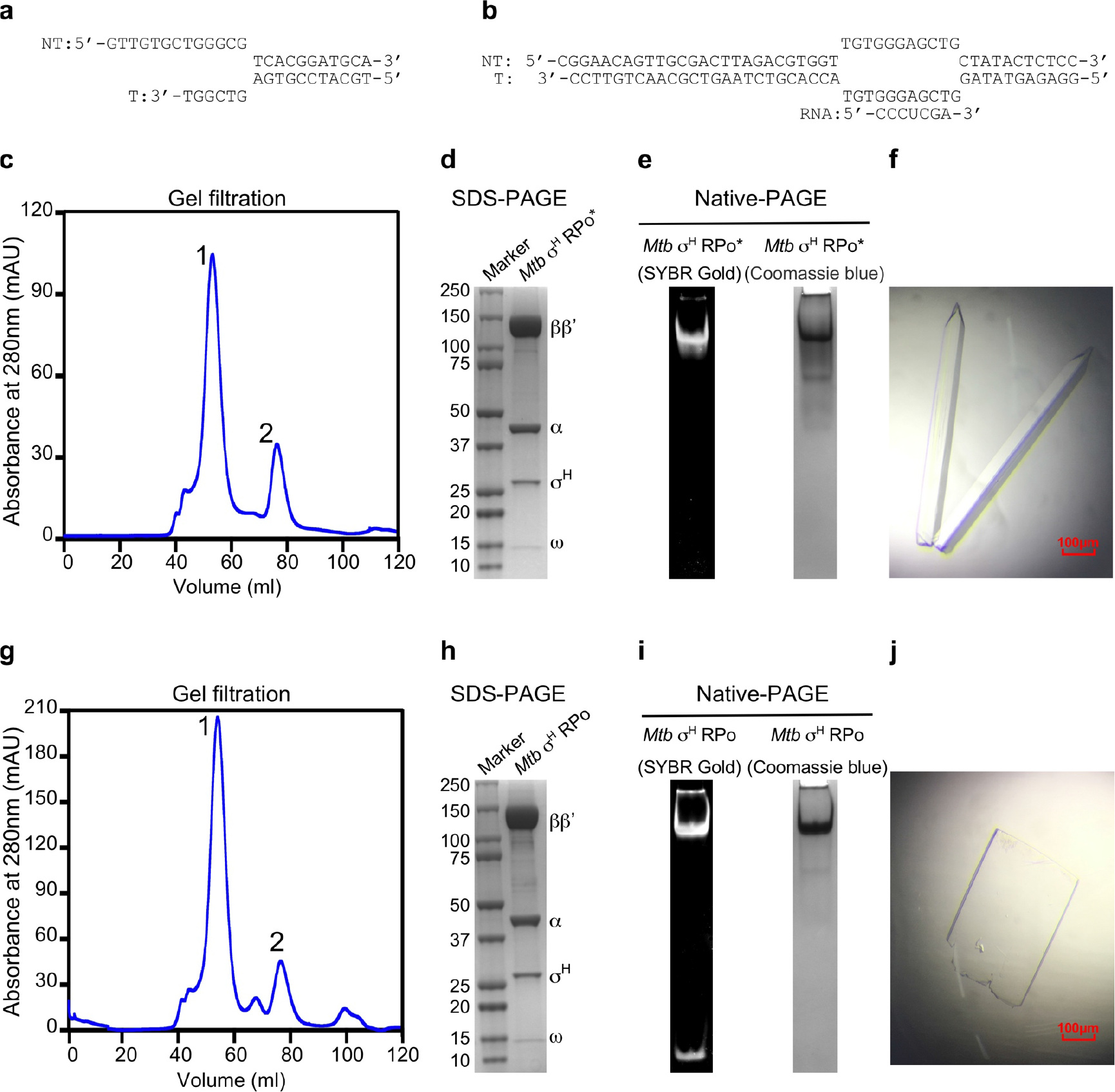
Protein and crystals preparation of *Mtb* σ^H^-RNAP holoenzyme and *Mtb* σ^H^-RPo. Related to Figure 1 and 3. **(a)** DNA scaffold for preparation of σ^H^-RPo* for crystallization of σ^H^-RNAP holoenzyme. **(b)** Nucleic-acid DNA scaffold for preparation of σ^H^-RPo. **(c)** Elution peaks of *Mtb* σ^H^-RPo* from a size-exclusion column. Peak 1 is the *Mtb* σ^H^-RPo* and peak 2 is the excess *Mtb* σ^H^. **(d)** The SDS-PAGE of peak 1 in (C). **(e)** The Native-PAGE of peak 1 in (C). The gel was first stained with SYBR Gold for nucleic acids and then with Coomassie blue for protein. **(f)** The crystals of *Mtb* σ^H^-holoenzyme. The DNA scaffold dissociated from RNAP during crystallization. **(g)** The elution peaks of *Mtb* σ^H^-RPo from a size-exclusion column. Peak 1 is the *Mtb* σ^H^-RPo and peak 2 is the excess *Mtb* σ^H^. **(h)** The SDS-PAGE of peak 1 in (G). **(i)** The Native-PAGE of peak 1 in (G). The gels were stained as above. **(j)** The crystal of *Mtb* σ^H^-RPo. * represents a RPo complex with a downstream fork promoter DNA.

**Supplementary Figure 2.**
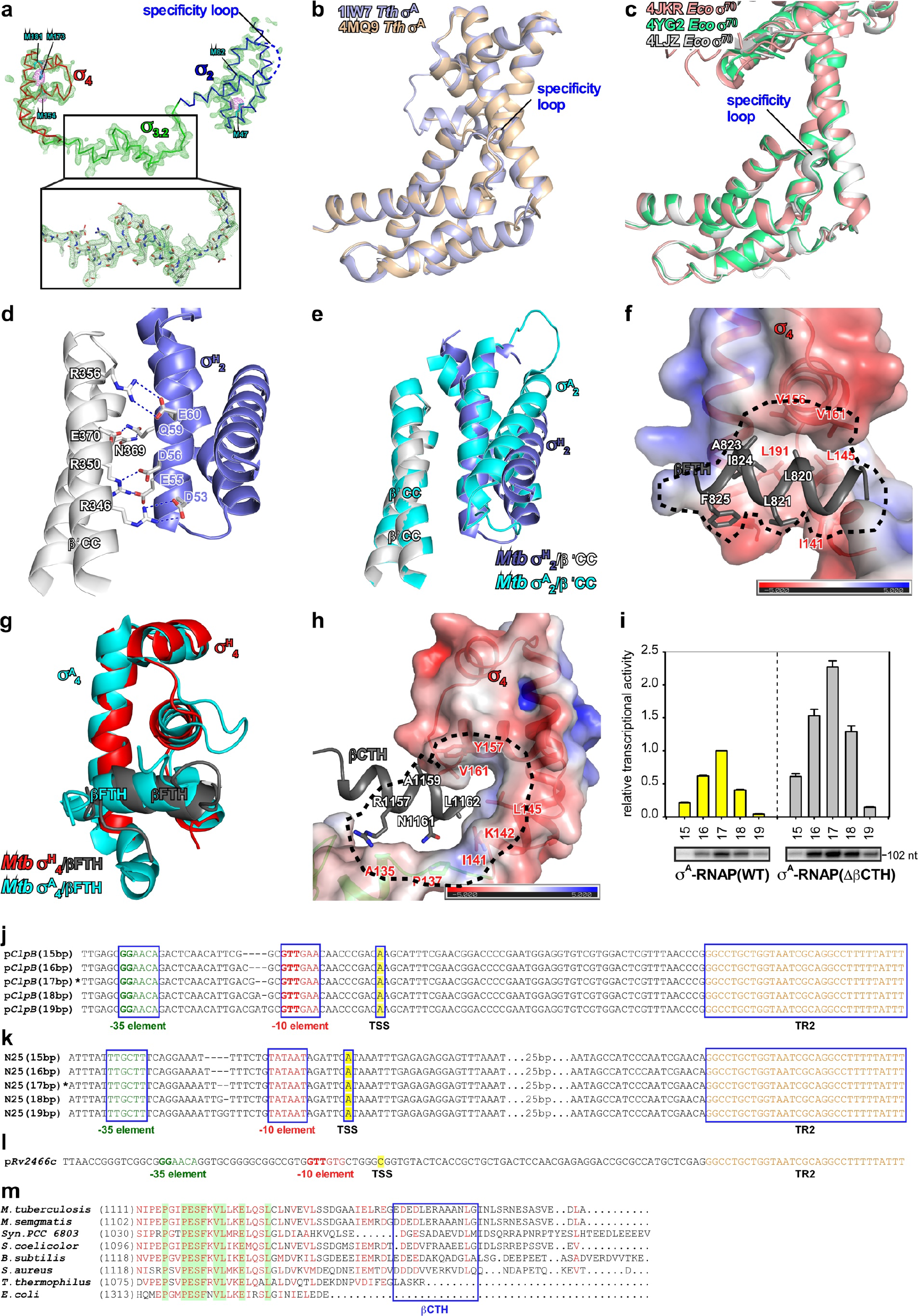
The detailed interaction between *Mtb* RNAP core enzyme and σ^H^. Related to Figure 1. **(a)** Electron density and model for σ^H^. Green mesh, simulated-annealing Fo-Fc difference map with σ^H^ omitted contoured at 2.5 σ Violet mesh, anomalous difference map contoured at 4 σ shows clear signals for 4 out of 5 Se atoms in σ^H^. σ^H^_2_, blue; σ^H^_3.2_, green; σ^H^_4_, red. **(b)** The specificity loops adopt ordered conformation in *T. thermophilus* σ^A^-RNAP holoenzymes (PDB: 1IW7 and 4MQ9) [https://www.rcsb.org/structure/1IW7; https://www.rcsb.org/structure/4MQ9]. **(c)** The specificity loops adopt ordered conformation in *E. coli* σ^70^-RNAP holoenzymes (PDB: 4JKR, 4YG2, and 4LJZ) [https://www.rcsb.org/structure/4JKR; https://www.rcsb.org/structure/4YG2; https://www.rcsb.org/structure/4LJZ]. **(d)** The interaction between σ^H^_2_ (blue) and RNAP-β′ coiled-coil (gray). H-bond, blue dash. **(e)** Superimposition of *Mtb* σ^H^_2_/β′CC (blue/gray) and *Mtb* σ^A^_2_/β′CC (cyan; PDB: 5UHA). **(f)** The interaction between σ^H^_4_ and RNAP βFTH. The electrostatic potential surface representation of σ^H^_4_ show a hydrophobic interface between σ^H^_4_ and RNAP βFTH. The electrostatic potential surface of σ^H^_4_ was generated using APBS tools in Pymol with partial changes determined by PDB2PQR server. Black dash, the hydrophobic groove; σ^H^_4_, red ribbon; RNAP βFTH, dark gray ribbon. **(g)** Superimposition of *Mtb* σ^H^_4_/βFTH (red/gray) and *Mtb* σ^A^_4_/βFTH (cyan; PDB: 5UHA). **(h)** The interaction between σ^H^_4_ and RNAP βCTH. The σ^H^_4_ electrostatic potential surface was generated as above. **(i)** The *in vitro* transcription activity of σ^A^-RNAP(WT) holoenzyme (yellow bars) or σ^A^-RNAP(ΔβCTH) holoenzymes (gray bars) from N25 promoter variants with −35/−10 spacer lengths of 15-19 base pairs. “102 nt” indicates length of terminated transcripts. The experiments were repeated for four times and the data were presented as mean ± S.E.M. **(j)** The promoter sequence of p*ClpB* derivatives for *in vitro* transcription activity of σ^H^-RNAP(WT) and σ^H^-RNAP(ΔβCTH). **(k)** The promoter sequence of N25 derivatives for *in vitro* transcription assay of σ^A^-RNAP(WT) and σ^A^-RNAP(ΔβCTH). **(l)** The promoter sequence of p*Rv2466c* for *in vitro* transcription assay of RNAP holoenzyme comprising σ^H^ derivatives. Source data of (I) are provided as a Source Data file. **(m)** a sequence alignment of RNAP βCTH of a few representative bacterial species.

**Supplementary Figure 3.**
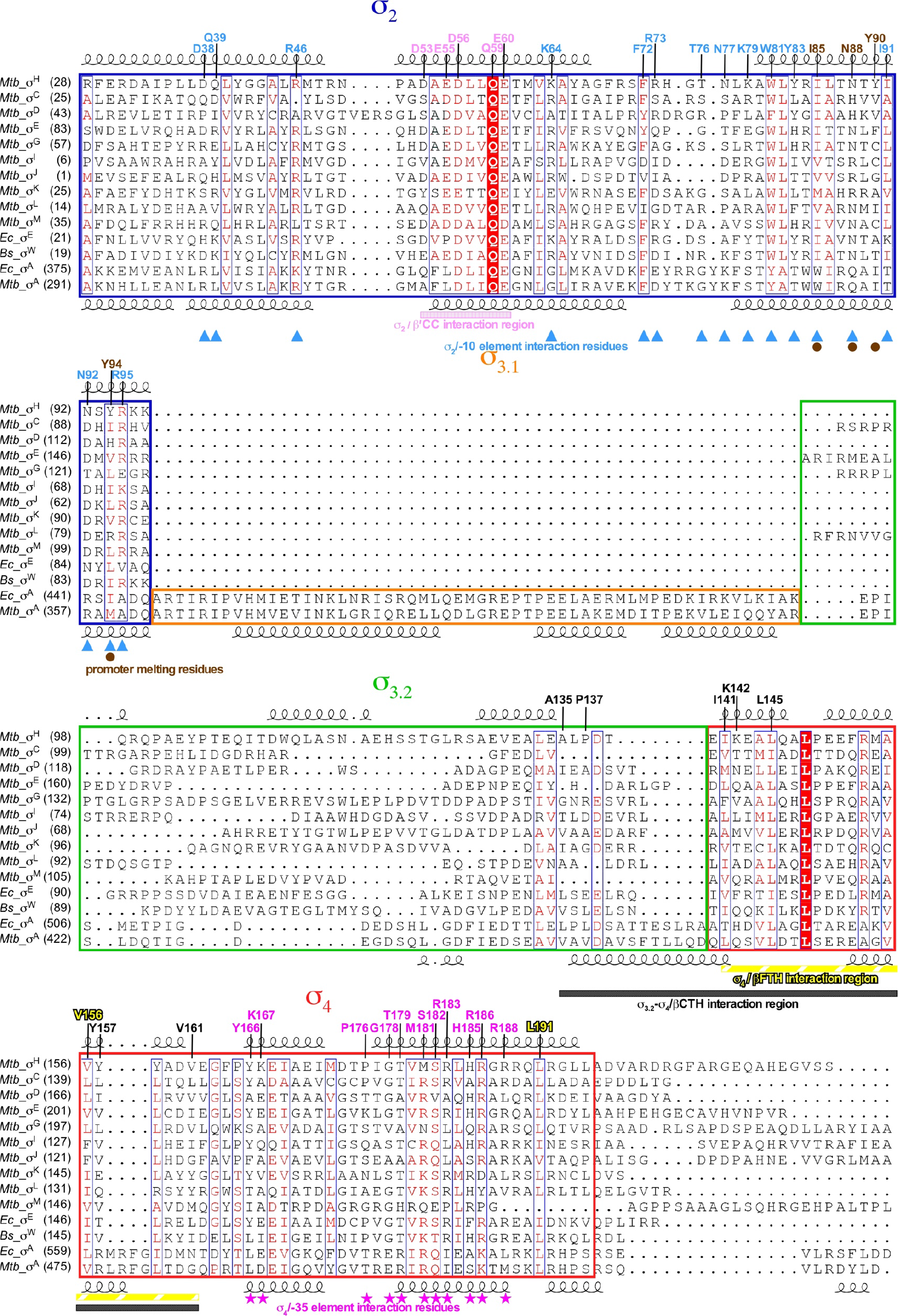
The sequence alignment of the ECF σ factors and primary σ^A^ factors. Related to Figures 1-5. The regions of σ_2_, σ_3.1_, σ_3.2_, and σ_4_ were indicated by blue, orange, green, and red boxes, respectively. Residues contacting the promoter −10 element are indicated by blue filled triangles; residues participating in promoter melting are indicated by brown filled circles; residues contacting the −35 elements are indicated by violet filled stars. The RNAP-β′CC interacting region, RNAP-βFTH interacting region, and RNAP-βCTH region on σ_2_ are highlighted by pink, yellow, and gray bars, respectively.

**Supplementary Figure 4.**
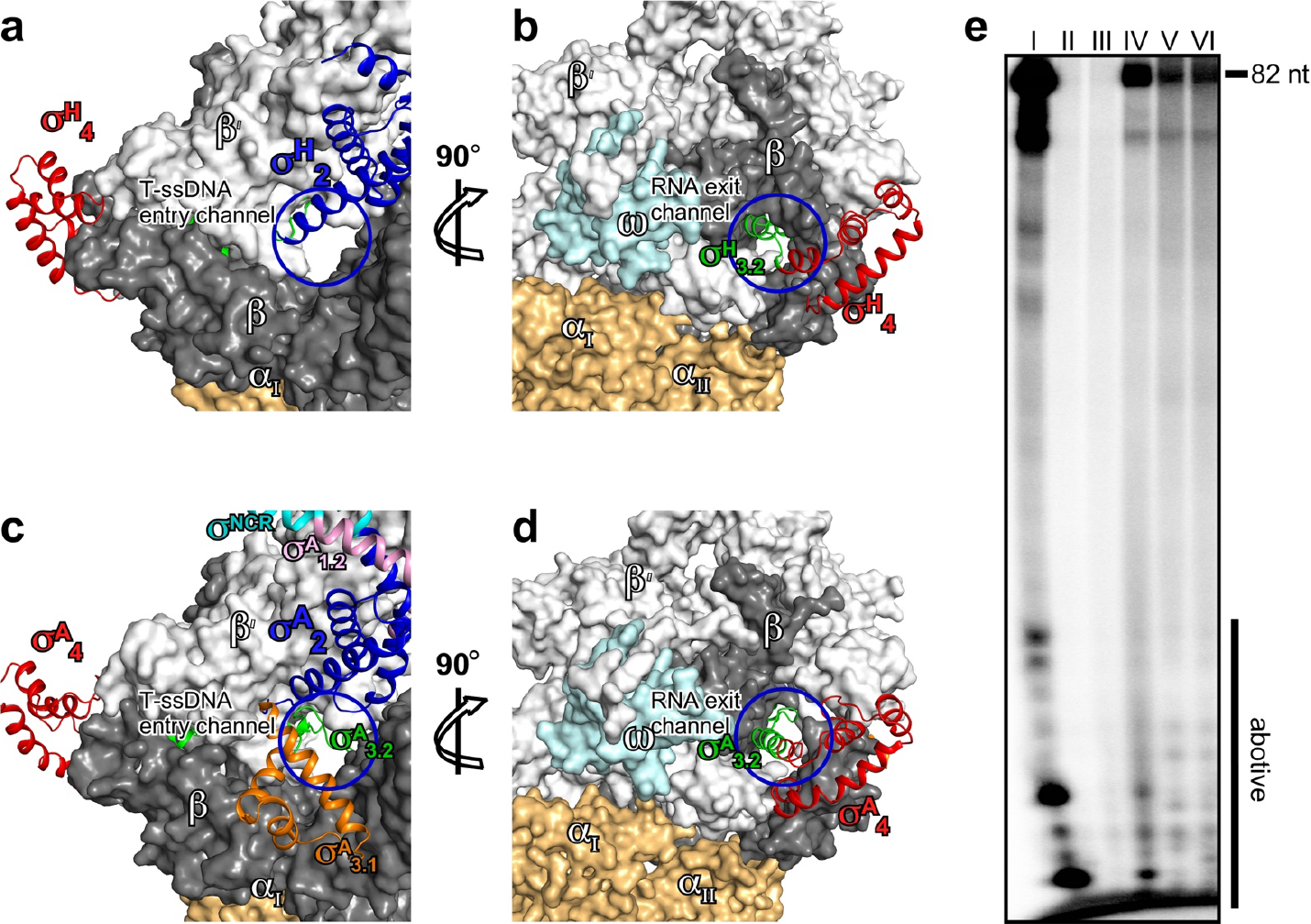
The σ_3.2_ pathway in *Mtb* σ^H^-RNAP and σ^A^-RNAP holoenzymes (extracted from PDB: 5UHA) [https://www.rcsb.org/structure/5UHA]. Related to Figure 2. **(a)** The entry channel of template single-strand DNA (T-ssDNA) created by σ^H^_2_ and σ^H^_3.2_. **(b)** The RNA exit channel blocked by σ^H^_3.2_. **(c)** The entry channel of template single-strand DNA (T-ssDNA) created by σ^A^_2_, σ^A^_3.1_, and σ^A^_3.2._ **(d)** The RNA exit channel blocked by σ^A^_3.2_. RNAP core enzyme is shown as surface and σ is shown as ribbon. Colors are as in main figures. **(e)** The *in vitro* transcription activity from p*Rv2466c* promoter of RNAP holoenzymes comprising σ^H^ derivatives. “82 nt” indicates length of terminated RNA transcripts. “abortive” indicates abortive transcripts. I, II, III, IV, V, VI indicate wild type σ^H^ or σ^H^ derivatives same as in Figure 2D. Source data of (E) are provided as a Source Data file.

**Supplementary Figure 5.**
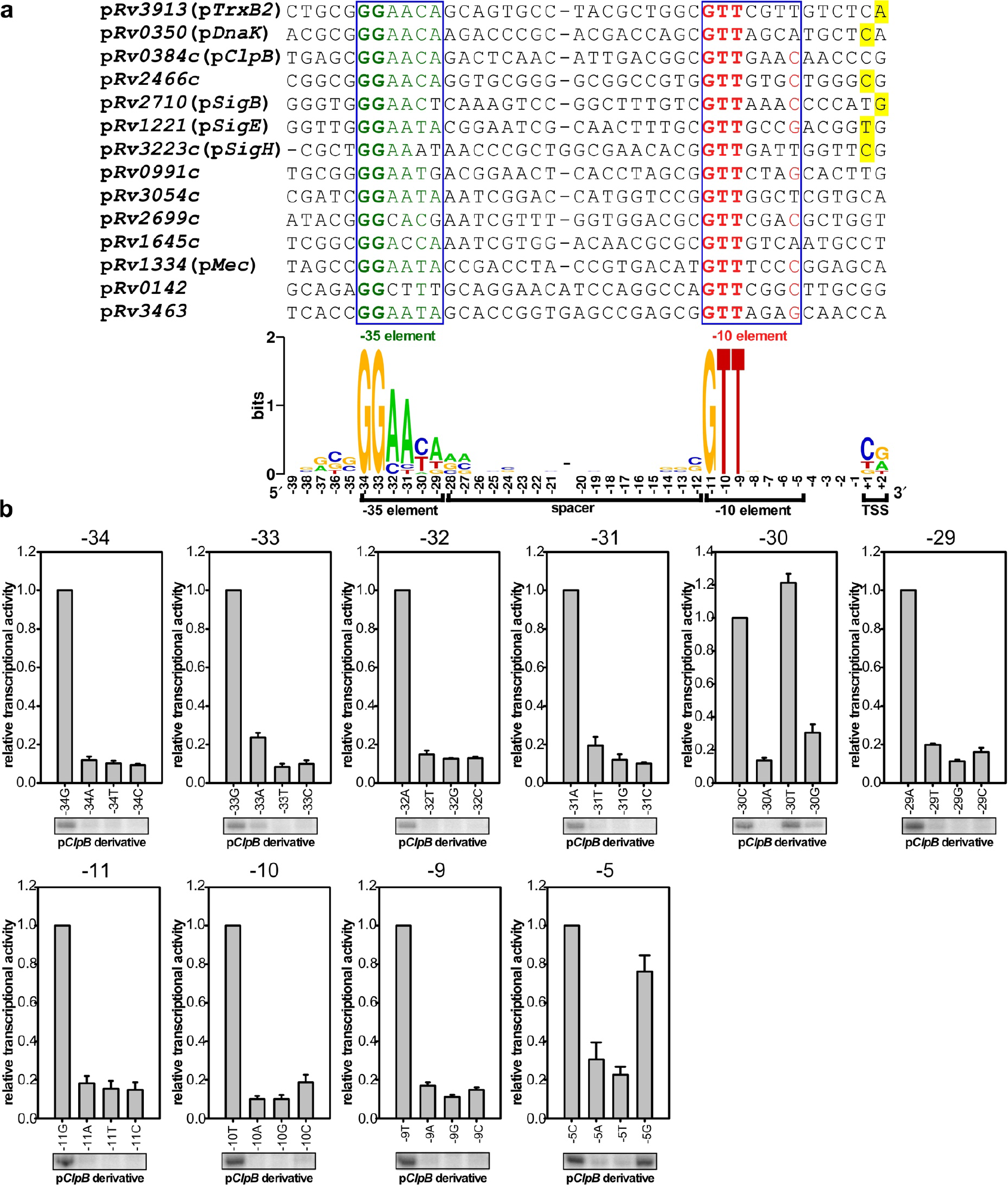
The consensus sequence of *Mtb* σ^H^-regulated promoters. Related to Figures 3-5. **(a)** The alignment of *Mtb* σ^H^-regulated promoter sequences. The sequence logo was generated on the WebLogo server. The conserved −35 element, green; the conserved −10 element, red; the transcription start site (TSS), yellow. **(b)** The *in vitro* transcription assays showing specificity of the promoter −35 and −10 elements. The representative run-off (122 nt) products were showed and quantified. The experiments were repeated for three times and the data were presented as mean ± S.E.M. Source data of (B) are provided as a Source Data file.

**Supplementary Figure 6.**
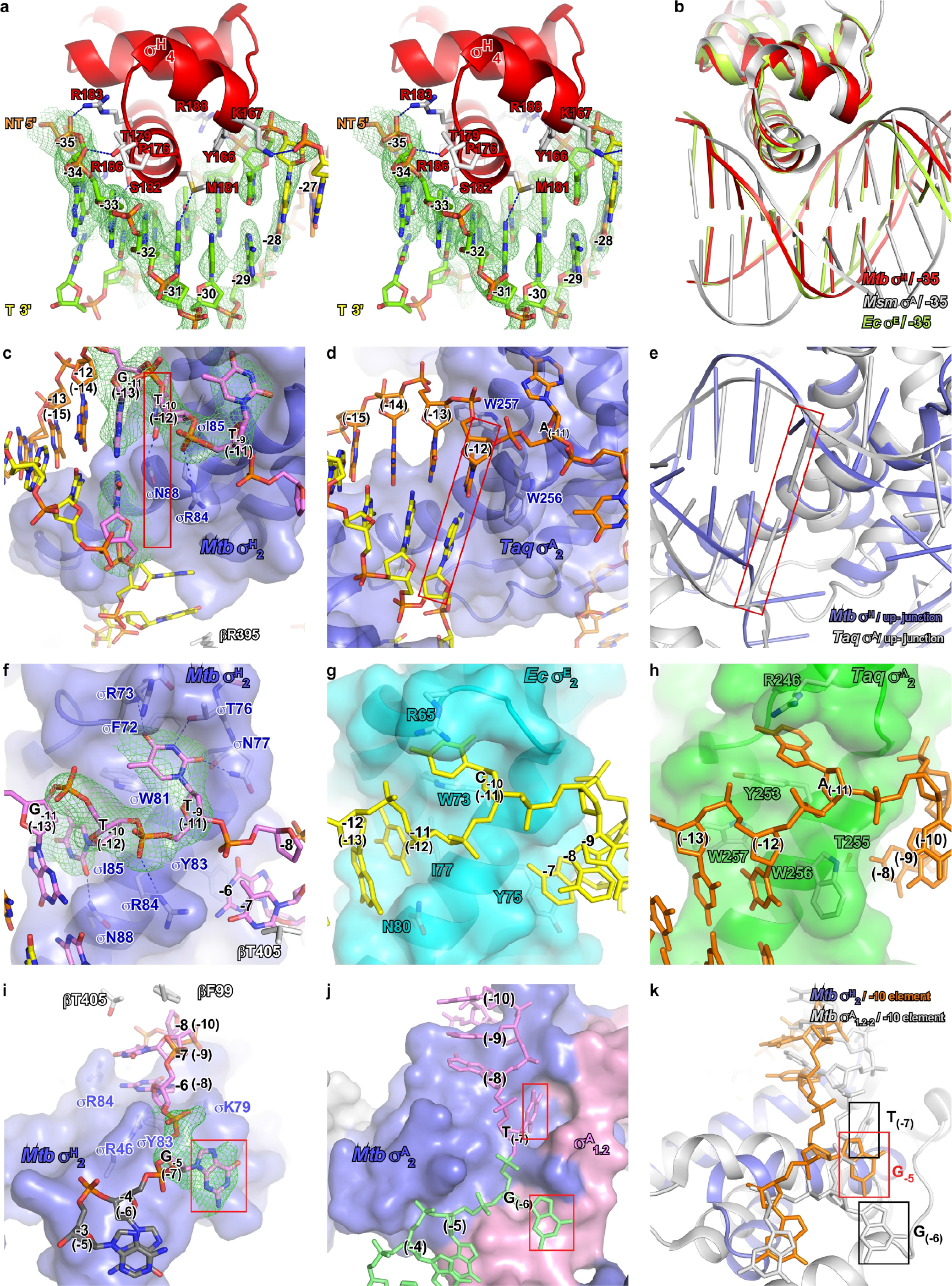
The detailed interactions of σ^H^-RNAP with promoter DNA −35/−10 elements and the comparison with σ^A^-RPo. Related to Figure 4. The *Mtb* σ^A^-RPo structure was chosen for superimposition with *Mtb* σ^H^-RPo unless the interactions to be compared is unavailable in *Mtb* σ^A^-RPo. **(a)** The stereo presentation of detail interactions between σ^H^_4_ and the −35 dsDNA. Color as in Figure 4A. **(b)** The superimposition of *Mtb* σ^H^_4_/−35 element (red), *Ec* σ^E^_4_/−35 element (green; PDB: 2H27) [https://www.rcsb.org/structure/2H27] and *Msm* σ^A^_4_/−35 element (gray; PDB: 5TW1) [https://www.rcsb.org/structure/5TW1] structures. **(c)** σ^H^_2_ melts promoter DNA at the junction of −11/−10 (corresponding to (−13)/(−12) of promoters for the primary σ factor). **(d)** σ^A^_2_ melts promoter DNA at the junction of (−12)/(−11) (PDB: 4XLN) [https://www.rcsb.org/structure/4XLN]. **(e)** The comparison of promoter melting by *Mtb* σ^H^-RNAP (blue) and *Taq* σ^A^-RNAP (gray; PDB: 4XLN). **(f)** *Mtb* σ^H^ inserts the bases of T_(−12)_(nt) and T_(−11)_ (nt) into pockets. **(g)** *Ec* σ^E^ inserts the bases of T_(−12)_(nt) and C_(−10)_(nt) into pockets (PDB: 4LUP) [https://www.rcsb.org/structure/4LUP]. **(h)** *Taq* σ^A^ inserts the base of A_(−11)_(NT) into a pocket (PDB: 4XLN). **(i)** Nucleotides at −7 and −6 positions of the −10 element ssDNA were stacked between βT405 and σY83 and G_−5_(nt) is inserted into a pocket in σ^H^-RPo. **(j)** Nucleotides at −10, −9 and −8 positions of the nontemplate DNA were stacked, T_(−7)_(nt) of the −10 element and G_(−6)_(nt) of the discriminator element are inserted into pockets in *Mtb* σ^A^-RPo (PDB: 5UHA) [https://www.rcsb.org/structure/5UHA]. **(k)** The G_−5_(nt) in σ^H^-RPo (blue/orange) and T_(−7)_(nt) in σ^A^-RPo (gray; PDB:5UHA) are the last positions of the −10 element, respectively, and located on similar positions on σ. Green mesh, simulated-annealing Fo-Fc difference map with nucleic acids omitted contoured at 2.5 σ.

**Supplementary Figure 7.**
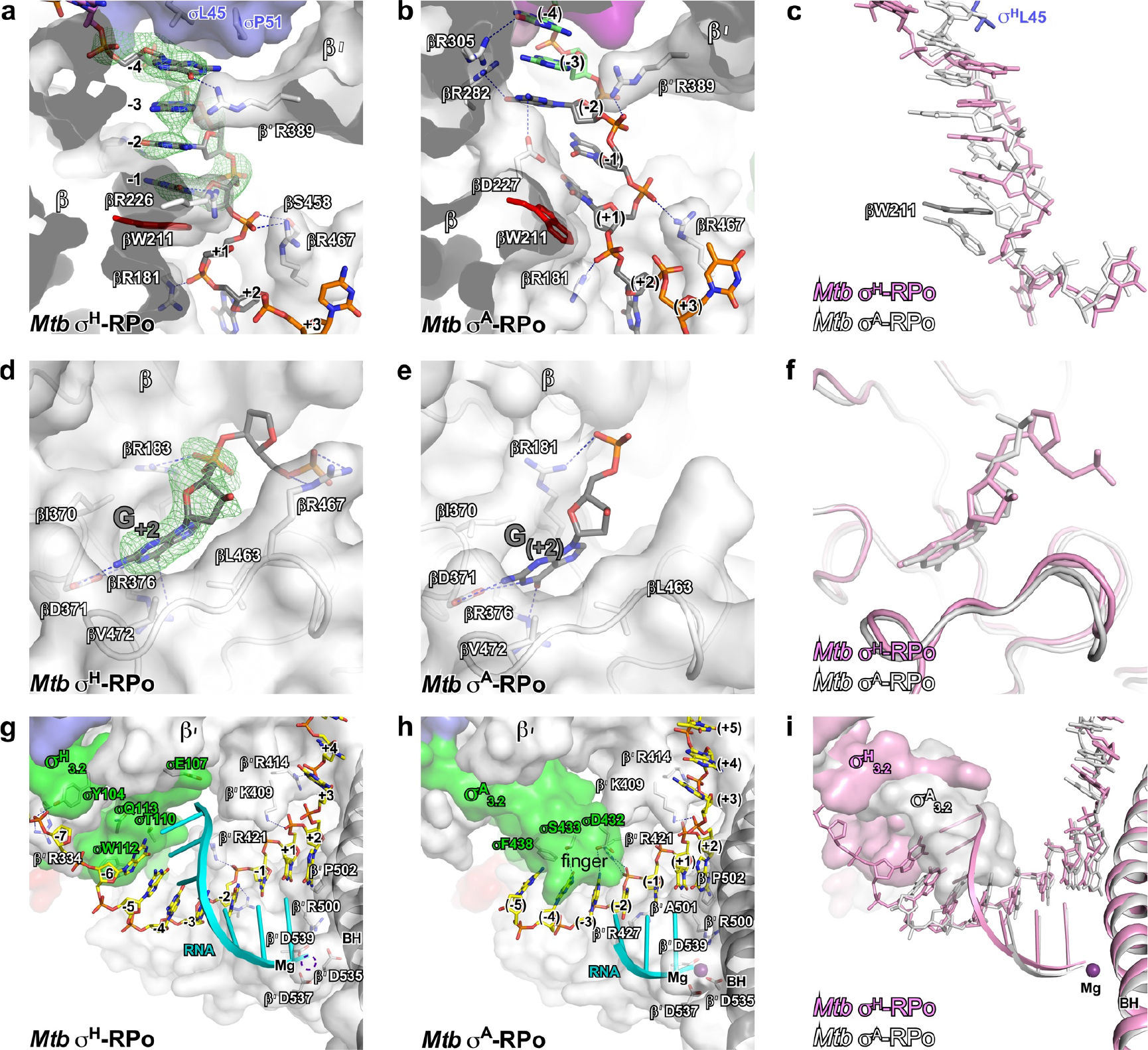
The detailed interaction of σ^H^-RNAP with CRE and DNA/RNA hybrid and the comparison with σ^A^-RPo. Related to Figure 5. **(a)** Nucleotides from −4 to −1 positions of nontemplate CRE were sandwiched between σL45 and βW211 in σ^H^-RPo. **(b)** The nucleotide from (−4) to (+1) positions of the nontemplate CRE were stacked by βW211 in *Mtb* σ^A^-RPo (PDB: 5UHA) [https://www.rcsb.org/structure/5UHA]. **(c)** The structure superimposition of the CRE elements in *Mtb* σ^H^-RPo (pink) and *Mtb* σ^A^-RPo (gray; PDB: 5UHA). **(d)** The G_+2_(nt) inserts into the ‘G’ pocket in σ^H^-RPo. **(e)** The G_(+2)_(nt) inserts into the ‘G’ pocket in *Mtb* σ^A^-RPo. **(f)** The interactions between the G_+2_(nt) and the ‘G’ pocket are the same in *Mtb* σ^H^-RPo (pink) and *Mtb* σ^A^-RPo (gray; PDB: 5UHA). **(g)** The interaction of the RNA/DNA hybrid with σ^H^_3.2_ and RNAP core enzyme in σ^H^-RPo. **(h)** The interaction of RNA/DNA hybrid with σ^A^_3.2_ and RNAP core enzyme in *Mtb* σ^A^-RPo (PDB: 5UHA). **(i)** The σ^A^_3.2_ in *Mtb* σ^A^-RPo (gray; PDB: 5UHA) reaches deeper into the active center compared with σ^H^_3.2_ in σ^H^-RPo (pink). Green mesh, simulated-annealing Fo-Fc difference map with nucleic acids omitted contoured at 2.5 σ.

## Reference

1. Browning DF, Busby SJ. Local and global regulation of transcription initiation in bacteria. Nat Rev Microbiol, (2016).

2. Lee DJ, Minchin SD, Busby SJ. Activating transcription in bacteria. Annual review of microbiology 66, 125–152 (2012).

3. Feklistov A, Darst SA. Structural basis for promoter-10 element recognition by the bacterial RNA polymerase sigma subunit. Cell 147, 1257–1269 (2011).

4. Feklistov A, Sharon BD, Darst SA, Gross CA. Bacterial sigma factors: a historical, structural, and genomic perspective. Annu Rev Microbiol 68, 357–376 (2014).

5. Ruff EF, Record MT, Jr., Artsimovitch I. Initial events in bacterial transcription initiation. Biomolecules 5, 1035–1062 (2015).

6. Saecker RM, Record MT, Jr., Dehaseth PL. Mechanism of bacterial transcription initiation: RNA polymerase - promoter binding, isomerization to initiation-competent open complexes, and initiation of RNA synthesis. J Mol Biol 412, 754–771 (2011).

7. Paget MS. Bacterial Sigma Factors and Anti-Sigma Factors: Structure, Function and Distribution. Biomolecules 5, 1245–1265 (2015).

8. Gruber TM, Gross CA. Multiple sigma subunits and the partitioning of bacterial transcription space. Annual review of microbiology 57, 441–466 (2003).

9. Mascher T. Signaling diversity and evolution of extracytoplasmic function (ECF) sigma factors. Curr Opin Microbiol 16, 148–155 (2013).

10. Staron A, Sofia HJ, Dietrich S, Ulrich LE, Liesegang H, Mascher T. The third pillar of bacterial signal transduction: classification of the extracytoplasmic function (ECF) sigma factor protein family. Mol Microbiol 74, 557–581 (2009).

11. Helmann JD. Bacillus subtilis extracytoplasmic function (ECF) sigma factors and defense of the cell envelope. Curr Opin Microbiol 30, 122–132 (2016).

12. Fiebig A, Herrou J, Willett J, Crosson S. General Stress Signaling in the Alphaproteobacteria. Annu Rev Genet 49, 603–625 (2015).

13. Woods EC, McBride SM. Regulation of antimicrobial resistance by extracytoplasmic function (ECF) sigma factors. Microbes Infect 19, 238–248 (2017).

14. Flentie K, Garner AL, Stallings CL. Mycobacterium tuberculosis Transcription Machinery: Ready To Respond to Host Attacks. J Bacteriol 198, 1360–1373 (2016).

15. Rodrigue S, Provvedi R, Jacques PE, Gaudreau L, Manganelli R. The sigma factors of Mycobacterium tuberculosis. FEMS Microbiol Rev 30, 926–941 (2006).

16. Sachdeva P, Misra R, Tyagi AK, Singh Y. The sigma factors of Mycobacterium tuberculosis: regulation of the regulators. FEBS J 277, 605–626 (2010).

17. Shultzaberger RK, Chen Z, Lewis KA, Schneider TD. Anatomy of Escherichia coli sigma70 promoters. Nucleic Acids Res 35, 771–788 (2007).

18. Yuzenkova Y, Tadigotla VR, Severinov K, Zenkin N. A new basal promoter element recognized by RNA polymerase core enzyme. Embo J 30, 3766–3775 (2011).

19. Barne KA, Bown JA, Busby SJ, Minchin SD. Region 2.5 of the Escherichia coli RNA polymerase sigma70 subunit is responsible for the recognition of the ‘extended-10’ motif at promoters. Embo J 16, 4034–4040 (1997).

20. Feklistov A, et al. A basal promoter element recognized by free RNA polymerase sigma subunit determines promoter recognition by RNA polymerase holoenzyme. Mol Cell 23, 97–107 (2006).

21. Campbell EA, et al. Structure of the bacterial RNA polymerase promoter specificity sigma subunit. Mol Cell 9, 527–539 (2002).

22. Bae B, Feklistov A, Lass-Napiorkowska A, Landick R, Darst SA. Structure of a bacterial RNA polymerase holoenzyme open promoter complex. eLife 4, (2015).

23. Feng Y, Zhang Y, Ebright RH. Structural basis of transcription activation. Science 352, 1330–1333 (2016).

24. Zhang Y, et al. Structural basis of transcription initiation. Science 338, 1076–1080 (2012).

25. Koo BM, Rhodius VA, Nonaka G, deHaseth PL, Gross CA. Reduced capacity of alternative sigmas to melt promoters ensures stringent promoter recognition. Genes Dev 23, 2426–2436 (2009).

26. Basu RS, et al. Structural basis of transcription initiation by bacterial RNA polymerase holoenzyme. J Biol Chem 289, 24549–24559 (2014).

27. Zuo Y, Steitz TA. Crystal structures of the E. coli transcription initiation complexes with a complete bubble. Mol Cell 58, 534–540 (2015).

28. Duchi D, et al. RNA Polymerase Pausing during Initial Transcription. Molecular cell 63, 939–950 (2016).

29. Petushkov I, Esyunina D, Mekler V, Severinov K, Pupov D, Kulbachinskiy A. Interplay between sigma region 3.2 and secondary channel factors during promoter escape by bacterial RNA polymerase. Biochem J 474, 4053–4064 (2017).

30. Samanta S, Martin CT. Insights into the mechanism of initial transcription in Escherichia coli RNA polymerase. J Biol Chem 288, 31993–32003 (2013).

31. Rhodius VA, et al. Design of orthogonal genetic switches based on a crosstalk map of sigmas, anti-sigmas, and promoters. Mol Syst Biol 9, 702 (2013).

32. Campagne S, Allain FH, Vorholt JA. Extra Cytoplasmic Function sigma factors, recent structural insights into promoter recognition and regulation. Curr Opin Struct Biol 30, 71–78 (2015).

33. Lane WJ, Darst SA. The structural basis for promoter −35 element recognition by the group IV sigma factors. PLoS Biol 4, e269 (2006).

34. Campagne S, Marsh ME, Capitani G, Vorholt JA, Allain FH. Structural basis for −10 promoter element melting by environmentally induced sigma factors. Nat Struct Mol Biol 21, 269–276 (2014).

35. Hubin EA, et al. Structure and function of the mycobacterial transcription initiation complex with the essential regulator RbpA. eLife 6, (2017).

36. Lin W, et al. Structural Basis of Mycobacterium tuberculosis Transcription and Transcription Inhibition. Molecular cell 66, 169–179 e168 (2017).

37. Degen D, et al. Transcription inhibition by the depsipeptide antibiotic salinamide A. eLife 3, e02451 (2014).

38. Manganelli R, Voskuil MI, Schoolnik GK, Dubnau E, Gomez M, Smith I. Role of the extracytoplasmic-function sigma factor sigma(H) in Mycobacterium tuberculosis global gene expression. Mol Microbiol 45, 365–374 (2002).

39. Chakraborty A, et al. Opening and closing of the bacterial RNA polymerase clamp. Science 337, 591–595 (2012).

40. Raman S, Song T, Puyang X, Bardarov S, Jacobs WR, Jr., Husson RN. The alternative sigma factor SigH regulates major components of oxidative and heat stress responses in Mycobacterium tuberculosis. J Bacteriol 183, 6119–6125 (2001).

41. Song T, Song SE, Raman S, Anaya M, Husson RN. Critical role of a single position in the −35 element for promoter recognition by Mycobacterium tuberculosis SigE and SigH. J Bacteriol 190, 2227–2230 (2008).

42. Kim KY, Park JK, Park S. In Streptomyces coelicolor SigR, methionine at the −35 element interacting region 4 confers the −31′-adenine base selectivity. Biochem Biophys Res Commun 470, 257–262 (2016).

43. Juang YL, Helmann JD. A promoter melting region in the primary sigma factor of Bacillus subtilis. Identification of functionally important aromatic amino acids. J Mol Biol 235, 1470–1488 (1994).

44. Fenton MS, Lee SJ, Gralla JD. Escherichia coli promoter opening and −10 recognition: mutational analysis of sigma70. Embo J 19, 1130–1137 (2000).

45. Campbell EA, et al. Crystal structure of Escherichia coli sigmaE with the cytoplasmic domain of its anti-sigma RseA. Molecular cell 11, 1067–1078 (2003).

46. Sainsbury S, Niesser J, Cramer P. Structure and function of the initially transcribing RNA polymerase II-TFIIB complex. Nature 493, 437–440 (2013).

47. Murakami KS, Masuda S, Darst SA. Structural basis of transcription initiation: RNA polymerase holoenzyme at 4 A resolution. Science 296, 1280–1284 (2002).

48. Kulbachinskiy A, Mustaev A. Region 3.2 of the sigma subunit contributes to the binding of the 3′-initiating nucleotide in the RNA polymerase active center and facilitates promoter clearance during initiation. J Biol Chem 281, 18273–18276 (2006).

49. Pupov D, Kuzin I, Bass I, Kulbachinskiy A. Distinct functions of the RNA polymerase sigma subunit region 3.2 in RNA priming and promoter escape. Nucleic Acids Res 42, 4494–4504 (2014).

50. Lerner E, et al. Backtracked and paused transcription initiation intermediate of Escherichia coli RNA polymerase. Proceedings of the National Academy of Sciences of the United States of America, (2016).

51. Liu X, Bushnell DA, Kornberg RD. Lock and key to transcription: sigma-DNA interaction. Cell 147, 1218–1219 (2011).

52. Vvedenskaya IO, et al. Massively Systematic Transcript End Readout, “MASTER”: Transcription Start Site Selection, Transcriptional Slippage, and Transcript Yields. Mol Cell 60, 953–965 (2015).

53. Heyduk E, Kuznedelov K, Severinov K, Heyduk T. A consensus adenine at position −11 of the nontemplate strand of bacterial promoter is important for nucleation of promoter melting. J Biol Chem 281, 12362–12369 (2006).

54. Lim HM, Lee HJ, Roy S, Adhya S. A “master” in base unpairing during isomerization of a promoter upon RNA polymerase binding. Proc Natl Acad Sci U S A 98, 14849–14852 (2001).

55. Tomsic M, Tsujikawa L, Panaghie G, Wang Y, Azok J, deHaseth PL. Different roles for basic and aromatic amino acids in conserved region 2 of Escherichia coli sigma(70) in the nucleation and maintenance of the single-stranded DNA bubble in open RNA polymerase-promoter complexes. J Biol Chem 276, 31891–31896 (2001).

56. Banerjee R, Rudra P, Prajapati RK, Sengupta S, Mukhopadhyay J. Optimization of recombinant Mycobacterium tuberculosis RNA polymerase expression and purification. Tuberculosis (Edinb) 94, 397–404 (2014).

57. Otwinowski Z, Minor W. [20] Processing of X-ray diffraction data collected in oscillation mode. Methods Enzymol 276, 307–326 (1997).

58. McCoy AJ, Grosse-Kunstleve RW, Adams PD, Winn MD, Storoni LC, Read RJ. Phaser crystallographic software. J Appl Crystallogr 40, 658–674 (2007).

59. Emsley P, Cowtan K. Coot: model-building tools for molecular graphics. Acta Crystallogr D Biol Crystallogr 60, 2126–2132 (2004).

60. Adams PD, et al. PHENIX: a comprehensive Python-based system for macromolecular structure solution. Acta Crystallogr D Biol Crystallogr 66, 213–221 (2010).

